# Antibodies, Memory B Cells, and Antigen Valency Reshape B Cell Responses to Drifted Influenza Virus Vaccination

**DOI:** 10.1101/2025.11.07.687210

**Authors:** Laura Reusch, Nimitha R. Mathew, Karin Schön, Ivan Kosik, James S. Gibbs, Madeleine C. Mankowski, Danica F. Besavilla, Jonathan W. Yewdell, Mats Bemark, Davide Angeletti

## Abstract

Antigenic drift in influenza A virus hemagglutinin (HA) limits humoral protective immunity. Here, we combine cell fate mapping with adoptive transfer of antigenic-site-specific antibodies (Abs) and memory B cells (MBCs) with moderately drifted HA vaccination in mice to better understand how this influences immune escape and protective responses. We demonstrate that drift in vaccine antigens affects MBC reactivation and naïve B cell responses in germinal centers (GC). Strikingly, passively transferred monoclonal and polyclonal Abs suppress cognate epitope-specific GC B cell responses only when the vaccine antigen was multivalent while responses to monovalent recombinant trimeric HA remain unaffected. Using MBC and Abs co-transfer we unveil that antigenic site-specific suppression is more potent in blocking MBC rather than naïve B cells entry into GC. In addition, we show that MBC hamper naïve B cell recruitment to GC even in the absence of antibody transfer through local differentiation and Ab release in the responding lymph node. Altogether, our study reveals that serum Ab feedback depends on vaccine valency, while pre-existing MBC alone without Abs present can reshape immunodominance of naïve B cells, with critical practical implications for rational universal influenza vaccine design.

## INTRODUCTION

Human influenza A viruses (IAV) exhibit rapid antigenic evolution in their hemagglutinin (HA) and neuraminidase (NA) virion surface glycoproteins. These “drifted” variants emerge to evade existing neutralizing antibodies (Abs) induced by prior infection and vaccination ^1, 2^. Often, a few mutations in critical sites of HA are sufficient to evade protection, with an average of 0.0011 mutations per antigenic site per year^3^. Memory B cells (MBC) are often better equipped than Abs to deal with drifted variants, owing to their wider B cell receptor (BCR) diversity than Ab-secreting plasma cells ^4, 5, 6^. Despite many studies, it has not been fully elucidated how drifted viruses and vaccines, pre-existing Abs, and MBC, interact to reshape humoral immunity at detailed epitope resolution. This is a central question in the quest for universal influenza vaccination ^7, 8^.

After infection/vaccination, MBC can arise both from the rapid extrafollicular (EF) response or the slower germinal center (GC) reaction ^6, 9, 10^. Antigen-specific GC B cells experience several rounds of BCR diversification and selection, ultimately differentiating into either plasma cells (PC) or MBC ^11^. After exiting the GC, MBC either circulate or reside in infected tissues: the respiratory tract in the case of IAV ^8^. Upon antigen re-encounter, MBC either enter a secondary GC or quickly differentiate into antibody-secreting cells (ASC) ^11, 12^. A better understanding of factors affecting MBC fate decision upon antigen rechallenge will facilitate designing viral vaccines that optimize MBC differentiation.

The fate of MBC during recall responses depends on many cell intrinsic factors, including expression of specific surface molecules, antibody heavy chain class, and BCR affinity dependent on somatic mutations introduced into the Ab variable region ^12, 13, 14, 15, 16, 17, 18, 19, 20^. Extrinsic factors, including the amount and specificity of pre-existing Abs and CD4 T cells as well as antigenic distance between the priming and challenging antigens, also contribute to MBC fate choice ^21, 22, 23, 24, 25^. For instance, greater antigenic distance and more memory CD4 T cells increased MBC reentry into secondary GC, while pre-existing Abs favored ASC generation. In addition, monoclonal Abs (mAbs) have been shown to influence *de novo* B cell responses by suppressing differentiation of B cells specific for the mAb’s cognate antigenic site and thereby instead promoting B cell responses towards non-cognate antigenic sites ^24, 25, 26, 27, 28, 29, 30^.

Most studies that have addressed these topics relied on model antigens with few antigenic epitopes, transfer of mAbs, knock-in BCR mice with fixed antigen specificity or a combination of them. How immune responses are affected in a more physiological setting where a drifted antigen is used to challenge a host with a preexisting polyclonal Ab and/or MBC has not been tested.

For all but unvaccinated children under three years of age ^31^, human IAV infection/vaccination confronts a polyclonal Ab and MBC repertoire that dictates B cell responses ^32^ ^33^. The majority of protective Abs are specific for HA ^29^. For the H1 subtype of HA, most Abs bind five overlapping canonical antigenic sites within the HA head (namely Sa, Sb, Ca1, Ca2 and Cb; Figure 1A) ^34, 35, 36^. These are targeted hierarchically after intranasal (i.n.) infection or vaccination, with the early response focusing on Cb and the later responses on Sb ^29^. Abs to non-canonical (n.c.) antigenic sites, including the receptor binding site (RBS) ^37^ or the stem ^38^, are usually cross-reactive between IAV strains but are immunosubdominant in animals and humans ^2, 29, 39, 40, 41, 42^. Intriguingly, several studies revealed a polyclonal Ab response highly focused on a single epitope, in approximately a fifth of individuals ^43, 44, 45, 46^: serum Ab recognition of HA in these people is easily abrogated by a single amino substitution in one of the canonical antigenic sites. It is easy to imagine scenarios where polyclonal Ab responses are more focused toward a single antigenic site while MBC target a wider array of epitopes, reflecting the repertoire diversity between PC and MBC ^4, 5, 6^. In this case, how will the recall response be influenced? Will MBC reactivation and naïve B cells recruitment into GC be impacted differently? And how will it contribute to the phenomenon known as original antigenic sin ^47^, the boosting of pre-existing antibodies weakly cross-reactive with the challenging immunogen?

**Figure 1.**
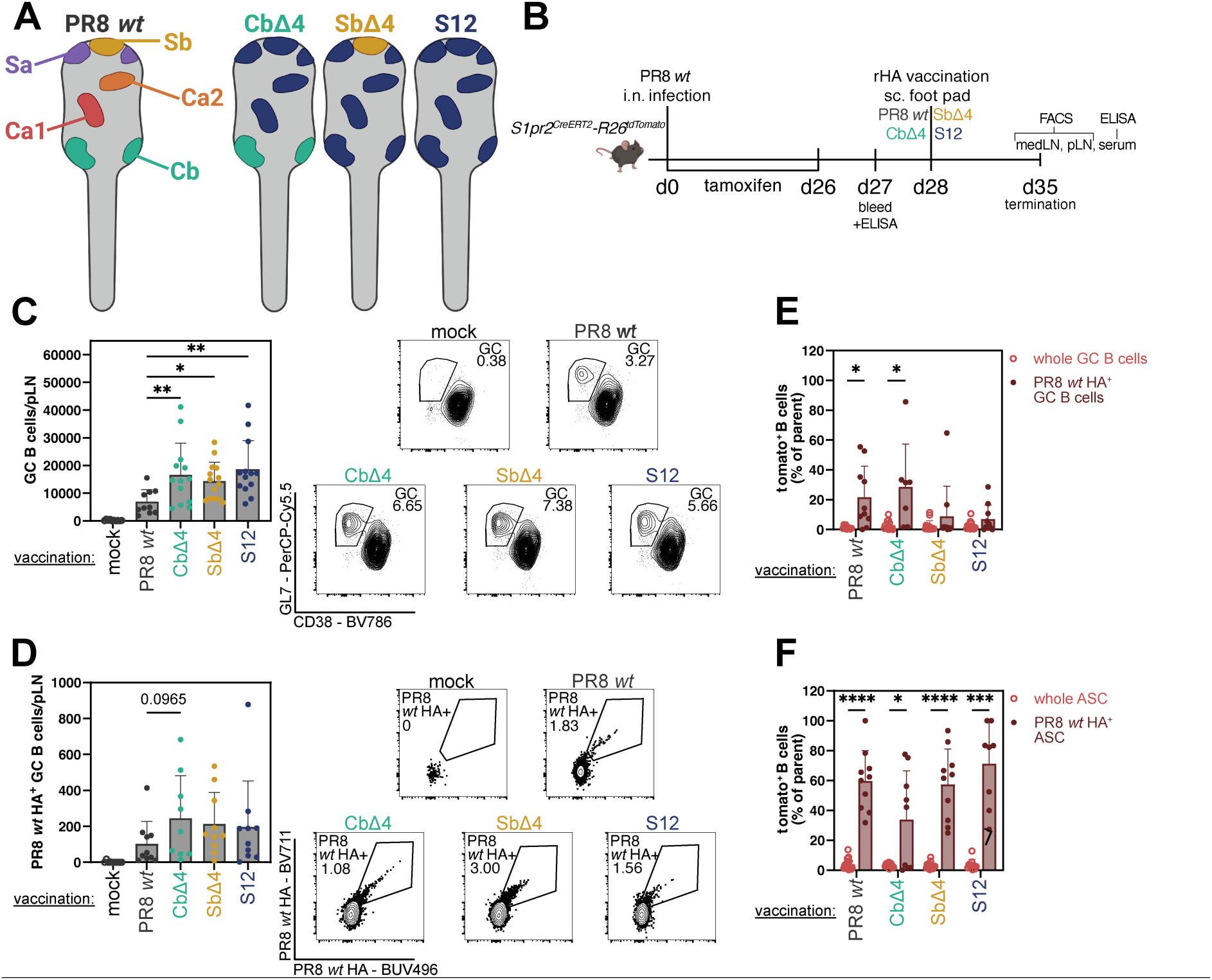
A drifted rHA challenge reshapes secondary GC responses in the LN draining the challenge site. **A**: Simplified graphic of *wt* HA with canonical antigenic sites and Δ4 escape mutants: Cb in green, Sb in yellow, Sa in purple, Ca1 in red, Ca2 in orange. Escaped/drifted sites are in dark blue and non-canonical sites in grey (rest of protein). **B:** Experimental setup of cell fate tracking using the S1pr2-Tomato mouse model. **C:** Total count of GC B cells in pLN at 7 days post vaccination (dpv). Challenge groups are listed from left to right by antigenic distance from *wt* (left). Representative flow plots of GC B cells (lin^-^ B220^+^ CD138^low/-^ CD19^+^ CD38^-^ GL7^+^) in pLN at 7dpv (right). **D:** Total count of *wt* HA^+^ GC B cells in pLN at 7dpv. (left). Representative flow plots of *wt* HA^+^ GC B cells (lin^-^ B220^+^ CD138^low/-^ CD19^+^ CD38^-^ GL7^+^ *wt*-BV711^+^ *wt*-BUV496^+^) in pLN at 7dpv (right). **E:** Frequency of fate mapped (S1pr2- TdTomato^+^) cells among GC B cells and **F:** ASC (lin^-^ B220^+^ CD138^high^). Light red = whole GC B cell or ASC, dark red = *wt* HA^+^ GC B cells or ASC. p-values were calculated by unpaired t tests (**E, F**) or one-way ANOVA with all groups compared to PR8 *wt* HA vaccination using (**C, D**). (∗p < 0.05; ∗∗p < 0.01; ∗∗∗∗p < 0.0001). Lowest p value indicated if no significance detected. Figures represent data from two experiments with 3-7 mice per challenge group per experiment. At least 2 experiments/group were performed. All data presented as mean ± SD. Animals without any detectable anti-*wt* HA IgG serum titer at -1dpv were excluded from data analysis.

Here, we use cell fate tracking models, adoptive transfers of polyclonal Abs and MBC, and a panel of drifted mutants of IAV A/Puerto Rico/8/1934 (PR8) HA to dissect how pre-existing humoral immunity redirects memory and *de novo* B cell responses upon IAV vaccination.

## RESULTS

### Antigen-specific MBC re-entry into secondary germinal centers is dependent on drift of vaccine antigen

To track the fate of antigenic site-specific cells, we used a panel of escape mutant viruses and corresponding HA based on the wildtype HA (*wt*) from the PR8 H1N1 influenza strain ^29^. Each “Δ4 mutant” expresses just one of the five original canonical antigenic sites (**Error! Reference source not found.**A). We focused on Cb and Sb antigenic sites as they represent early and late immunodominant sites after infection that are spatially oriented to minimize steric hindrance of simultaneous Ab binding ^29^. In addition, we used S12 as a fully drifted antigen that is poorly recognized by anti-*wt* HA sera ^48^. The mutants used in this work represent a timeframe of two to four influenza seasons ^49^, corresponding to an antigenic drift of up to 2.1% or ≤12 amino acids (aa) (**Error! Reference source not found.**A) ^29^, a likely time frame for an individual to be reinfected. These represent an improvement of the Δ4 panel used in one of our prior studies ^29^, as we have optimized the panel to abolish residual Ab binding to a cryptic epitope present in the Sb site defined by residues 189 and 198 (H1 numbering)^50^.

We first investigated the ability of drifted vaccines to reactivate antigenic-site specific MBC elicited by a prior infection. We fate mapped GC B cells and their progeny generated during the primary infection using *S1pr2^CreERT^*^2^*-R26^tdTomato^*mice (S1pr2-Tomato) ^51^ infected i.n. with *wt* virus and treated with tamoxifen every second day after infection until rechallenge, to irreversibly label GC B cells and their progeny (Figure 1B). At 28dpi, mice were vaccinated with 10 μg recombinant HA vaccine (rHA) in the left footpad, thereby anatomically separating the draining lymph nodes (dLN) targeted by the primary infection (the mediastinal lymph node, medLN) and the secondary vaccination (the popliteal lymph node, pLN) (Figure 1B). The vaccine antigens used were either *wt*, CbΔ4, SbΔ4 or S12. Importantly, previous work has primarily investigated prime-boost vaccination but rarely infection-induced priming, which is physiologically relevant in most individuals ^21, 22, 23^.

Despite the relatively small changes between *wt* and Δ4 HAs, the total number of GC B cells in the pLN was significantly lower after a homologous *wt* rHA challenge than Δ4 or S12 rHA challenges (Figure 1C; Supplementary Figure 1B). This is consistent with previous prime and boost studies, where homologous vaccination generates small secondary GCs and low MBC re-entry ^21, 22, 23^. The increased GC response correlated with a greater total number but not frequency of HA *wt*-binding (*wt* HA^+^) B cells in the response to drifted *vs. wt* rHA antigen (Figure 1D; Supplementary Figure 1D), even though the analysis excluded naïve B cells responding to drifted antigenic sites. Most *wt* HA^+^ cells exhibited a GC phenotype, especially when mice were vaccinated using a drifted rHA (Supplementary Figure 1E), suggesting some level of Ab-mediated feedback when antigens were more similar. While total Tomato^+^ fate mapped MBC represented only 1-2% of vaccination-induced secondary GC B cells, in agreement with previous results ^21^ (Figure 1E and Supplementary Figure 1 F), among *wt* HA^+^ GC B cells the frequency of previously generated MBC re-entering the GC was much higher, depending on vaccine antigen (Figure 1E and Supplementary Figure 1F).

These findings indicate that heterologous vaccinations with antigenically distant antigens (SbΔ4 and S12) are better at inducing *de novo* responses, while homologous and less drifted vaccines (*wt* PR8 and CbΔ4) are more efficient in recalling infection-induced MBC to secondary GCs. As expected, when analyzing ASC, the majority of *wt* HA^+^ plasmablasts within the pLN were fate-mapped, regardless of challenging antigen (Figure 1F; Supplementary Figure 1F), in accordance with a model where MBC can rapidly differentiate to ASC to produce Abs ^22^.

### Vaccine antigen dictates the immunodominance pattern of recalled and *de novo* B cell responses

To determine the antigenic site specificity of responding B cells, we employed simultaneous Δ4 rHA staining with each probe bearing a different fluorophore (Figure 2A and Supplementary Figure 1B). After a multistep gating strategy to identify the B cell subpopulation of interest (Supplementary Figure 1B), two *wt* HA probes, conjugated to two distinct fluorochromes were used to minimize spurious binding. Next, binding to S12 HA defined specificity for non-canonical (n.c.) antigenic sites (grey). Among the *wt* HA^+^ S12^-^ B cells we then defined specific antigenic-site binders to Cb (green) and Sb (gold) by double staining with CbΔ4 HA and SbΔ4 HA probes. All *wt* HA^+^ cells that did not bind to S12 HA, CbΔ4 HA and SbΔ4 HA were defined as “others” (purple) binding to Ca1, Ca2 or Sa (Figure 2A and Supplementary Figure 1B). Alternatively, to define *de novo* reactivity to drifted sites, present only in the new variants (blue), we directly compared *wt* together with the drifted HA immediately after B cell gating: e.g. comparing the binding to *wt* HA and SbΔ4 HA with “new”, drifted reactivity defined as *wt* HA^-^ and SbΔ4 HA^+^ B cells (Figure 2B).

**Figure 2.**
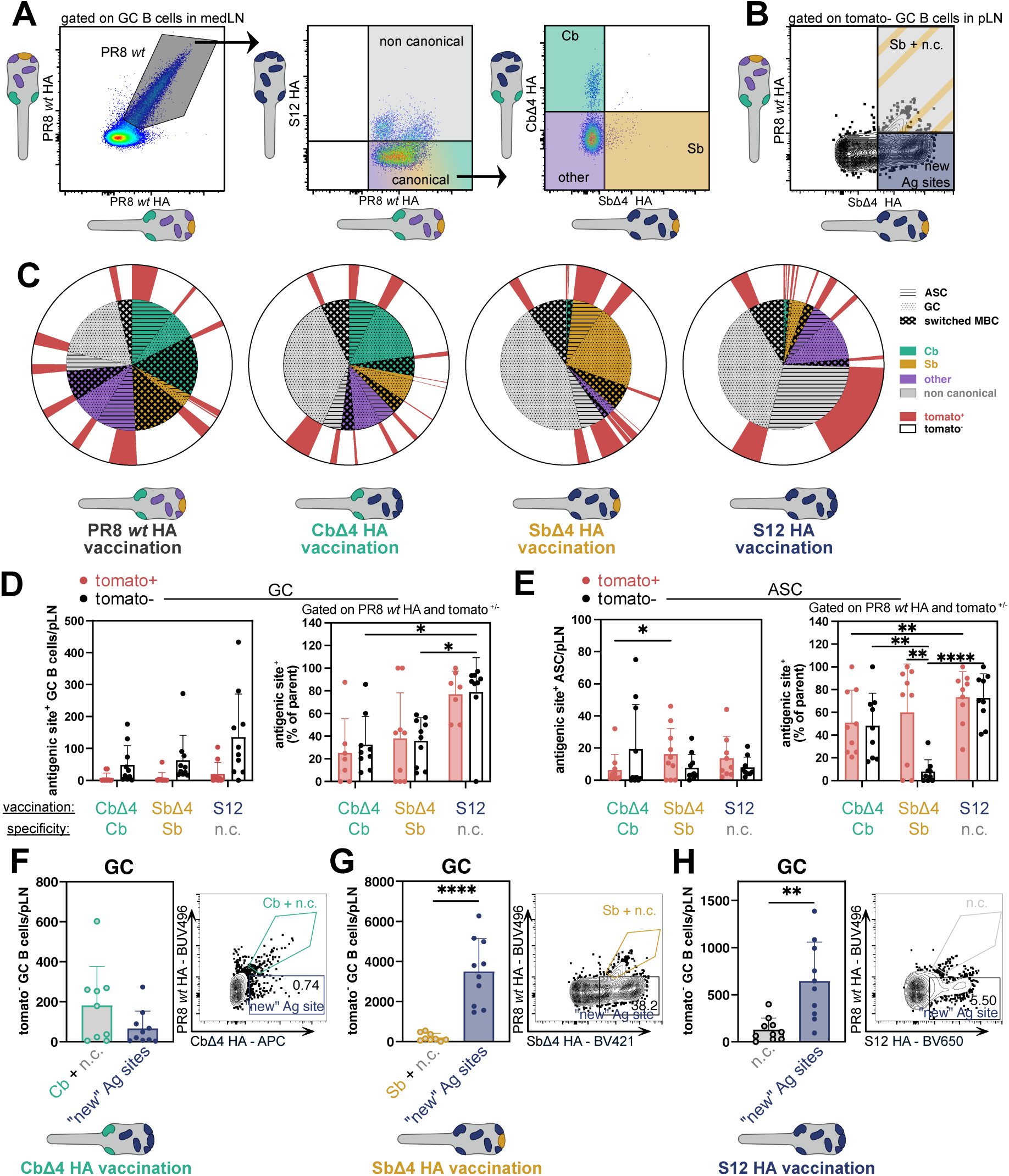
A drifted rHA challenge reshapes immunodominance patterns in draining LN. **A:** Gating strategy for antigenic site specificity that start from *wt* HA^+^ GC B cells to define non-canonical (S12 HA^+^), Cb^+^ (S12 HA^-^ SbΔ4 HA^-^ CbΔ4 HA^+^), Sb^+^ (S12 HA^-^ CbΔ4 HA^-^ SbΔ4 HA^+^), other^+^ (S12 HA^-^ CbΔ4 HA^-^ SbΔ4 HA^-^). Shown is a sample from medLN, 7dpv after *wt* rHA challenge in foot hock. **B:** Gating strategy for antigenic site specificity of “new” antigenic sites (dark blue) with example for SbΔ4 stainining. The same gating was used for CbΔ4 and S12 binding. It start from total GC B cells or ASC, and “new” antigenic sites^+^ (*wt* PR8 HA^-^ SbΔ4 or CbΔ4 or S12 HA^+^) and, respectively, Sb or Cb and/or n.c.+ (*wt* PR8 HA^+^ SbΔ4 or CbΔ4 or S12 HA^+^) **C:** Sunburst plots at 7 days after vaccination (dpv) of *wt* HA^+^cells divided by B cell phenotype (lined=ASC, dotted=GC, switched MBC=crossed lines), antigenic site specificity (Cb=green, Sb=yellow, other=purple, n.c. = grey) and fate mapping (white=not fate mapped, red=fate mapped/Tomato^+^) in pLN. Shown are plots for homologous/*wt* challenge as well as CbΔ4, SbΔ4 and S12 challenge. **D:** Total count and % per all *wt*-HA^+^ of GC B cells and **E:** ASC of challenge homologous antigenic site. Data is for Cb^+^ for CbΔ4 challenge, Sb^+^ for SbΔ4 challenge and n.c.^+^ for S12 challenge. Fate mapped indicated in red, non-fate mapped in black. **F:** Tomato^-^ Cb^+^ + n.c.^+^ and “new” antigenic site^+^ (*wt* HA^-^ CbΔ4 HA^+^) with representative plot from pLN at 7dpv, **G:** Sb^+^ + n.c.^+^ and “new” antigenic site ^+^ (*wt* HA^-^ SbΔ4 HA^+^) with representative plot from pLN at 7dpv, **H:** n.c.^+^ and “new” antigenic site ^+^ (*wt* HA^-^ S12 HA^+^) with representative plot from pLN at 7dpv of GC B cells total count per pLN. p- values were calculated by unpaired t test (**F, G, H**) or two-way ANOVA (**D, E**). (∗p < 0.05; ∗∗p < 0.01; ∗∗∗∗p < 0.0001). Figures represent data from two experiments with 3-7 mice per challenge group per experiment. All data as mean ± SD. Animals without any anti-*wt* HA IgG serum titers at -1dpv were excluded from data analysis.

We first determined how vaccination with the drifted immunogens broadens the response towards the antigen. We thus divided all *wt* HA^+^ cells according to antigenic site specificity (Figure 2C, Supplementary Figure 2A-B). *wt* HA challenge generated relatively similar ASC, MBC and GC responses equally distributed across the four antigenic groups (Cb, Sb, other immunodominant sites and n.c.; Figure 2C). A drifted rHA challenge favored responses not only to the conserved antigenic site but also increased the relative targeting of n.c. sites (grey) which are also conserved between the prime and boost antigens (Figure 2C), but, importantly, poorly immunogenic in primary immune responses ^29, 50^. This applied to B cells derived from both fate-mapped recalled MBC (Figure 2C and Supplementary Figure 2A) and naïve B cells (Figure 2C and Supplementary Figure 2B). The majority of these specific B cells exhibited a GC phenotype suggesting ongoing clonal diversification (Supplementary Figure 2C). These data support the concept that drifted immunogen boosting in a polyclonal setting extends B cell responses to clones not involved in the primary response, shifting B cell immunodominance towards sites less targeted in the primary Ab response.

By cell numbers, GC responses to the boosted antigenic site (*i.e.* Cb for CbΔ4 boost, Sb for SbΔ4 boost and n.c. for S12 boost) were dominated by a *de novo* response and not re-entering MBC (Figure 2D). However, the frequency of antigenic-site specific GC B cells was similar across Tomato^+^ and Tomato^-^ populations and increased according to antigenic distance to the *wt* rHA prime (Figure 2D). The significance of this finding is that the boosted antigen site is immunodominant both among the naïve- and the MBC-derived GC responses, with the *de novo* responses dominating.

ASC generation demonstrated a more complex pattern. Total cell numbers of fate-mapped and naïve B cell were similar in number after SbΔ4 and S12 but not CbΔ4 rHA vaccination groups (Figure 2E). More fate mapped MBC became ASC after vaccination with SbΔ4 rHA compared to CbΔ4 rHA (Figure 2E). Further, after SbΔ4 rHA vaccination, the fraction of Sb-specific ASC was much higher in Tomato^+^ *vs.* Tomato^-^ ASC indicating that only recalled MBC could become Sb-specific ASCs, at this early time point (Figure 2E). This was not the case for Cb, similar to what was observed upon primary PR8 infection ^29^. The results so far showed that minor antigenic differences can lead to distinctive outcomes in MBC reactivation.

Vaccination with variant rHA introduces drifted antigenic sites (blue in Figure 1A, 2B and 2F- H). As the drifted sites are not present on *wt* HA, only naïve B cells are activated to respond to them. For CbΔ4 rHA vaccination, GC response was similar between “old” and “new” antigenic sites, but the ASC response almost exclusively targeted “old” antigenic sites (Figure 2F and Supplementary Figure 2D). Conversely, after either SbΔ4 rHA and S12 rHA vaccination, GC responses were dominated by cells specific for drifted sites (Figure 2G-H). For ASC the response changed depending on vaccine (Supplementary Figure 2D-F). As the Cb site is immunodominant early after primary infection ^29^ it is reasonable to speculate that for SbΔ4 and S12 rHA vaccinations, most of the GC response to the drifted antigenic sites would target the “new”, drifted Cb site. This differential responses to drifted antigenic sites may contribute to the variable prevalence of original antigenic sin in previous studies ^47^.

We next used cell fate tracking to explore how peripheral rHA vaccination alters ongoing infection induced medLN GC reactions. Footpad vaccination with drifted rHA but not *wt* rHA invigorated total GC B cell responses in lung draining medLN (Supplementary Figure 3A), including *wt* HA specific responses (Supplementary Figure 3B). Interestingly, SbΔ4 rHA and S12 rHA vaccination increased numbers of n.c. and drifted-epitope specific GC B cells in comparison to CbΔ4 rHA or *wt* rHA challenges (Supplementary Figure 3C) and led to higher number of ASC in medLN (Supplementary Figure 3D), including *wt* HA^+^ specific ASC (Supplementary Figure 3E-F).

Overall, our fine epitope mapping of B cell specificities demonstrates that recall and *de novo* responses target similar antigenic sites. However, the ability of targeting “new”, drifted antigenic sites was also on vaccine drift. These differences may be caused by relative exposure of the conserved antigenic sites, its immunodominance and/or by relative abundance of epitope-specific, pre-existing MBC and Abs.

### Antigenic-site specific boosting of Ab responses varies according to vaccine antigen

Which factors exactly do reshape the specificity of rHA boost-induced B cell responses? A likely suspect is serum IgG, previously shown to modify B cell responses ^22, 23, 24, 25, 30, 52^. To assess the HA antigenic site specificities of anti-HA IgG, we measured IgG titers against *wt*, CbΔ4, SbΔ4 and S12 rHA 1 day pre- and 7 days post vaccination (dpv) (Supplementary Figure 4A-D) form the same experiment (Figure 1B), plotting the area under the curve (AUC) differences (Figure 3A-D).

**Figure 3:**
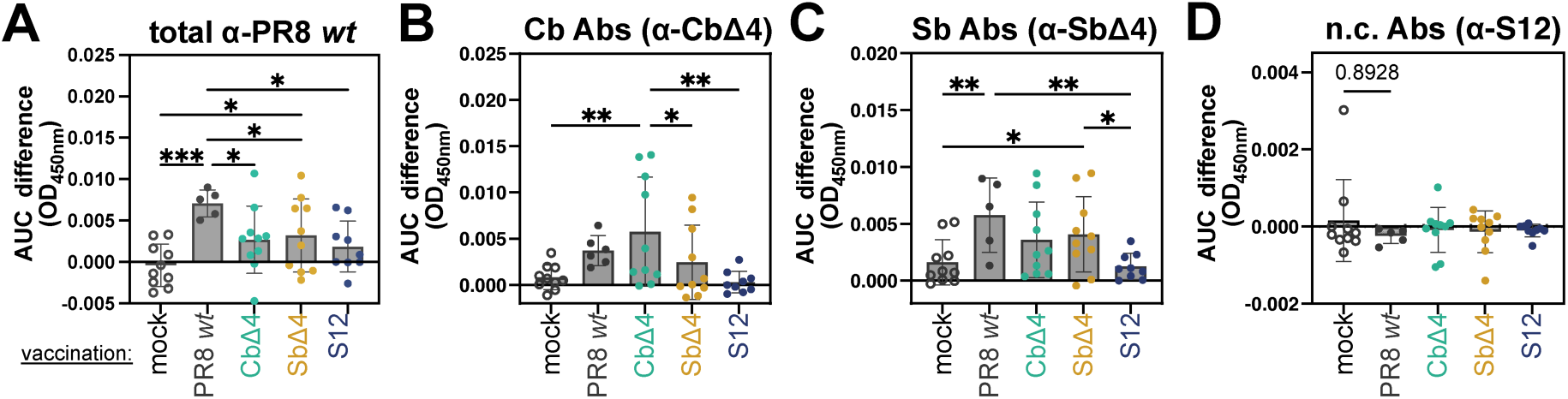
Immunodominance of serum Abs is influenced by drift in vaccine antigen. AUC of OD450nm difference between day -1 and 7 days post vaccination of IgG from serum. Plates, from left to right, were coated with rHA: **A:** *wt*, **B:** CbΔ4, **C:** SbΔ4, **D:** S12. p-values were calculated by one-way ANOVA, with all groups compared to *wt* HA challenge (homologous) or mock challenge (**A-F**). (∗p < 0.05; ∗∗p < 0.01; ∗∗∗∗p < 0.0001). Lowest p-value indicated if no significance detected. Figures represent data from two experiments with 3-7 mice per challenge group per experiment. At least two experiments per group were performed. All data as mean ± SD. Animals without any anti-*wt* HA IgG serum titers at -1dpv were excluded from data analysis.

Relative to mock vaccination, each vaccination condition enhanced *wt* HA specific IgG titers with homologous challenge showing the greatest increase, as expected (Figure 3A). A similar pattern was observed for Cb-specific Abs upon CbΔ4 rHA vaccination but not for Sb-specific Abs upon SbΔ4 rHA vaccination, consistent with the fate-mapped ASC data (Figure 3B-C and Supplementary Figure 4C-D). Intriguingly, n.c.-specific Ab titers did not increase with any rHA boost, contrary to the n.c.-specific GC B cell dominance observed after CbΔ4 or SbΔ4 vaccination and among ASC after S12 vaccination (Figure 3C; Supplementary Figure 4A-C). This suggests that n.c.-specific B cells remain in GCs, preferentially become MBC or ASC at later time point after vaccination. Alternatively, ASC may secrete low avidity Abs not detected by our standard ELISA conditions. These findings demonstrate that different heterologous vaccinations alter the balance of antigenic site-specific Abs, with site-specific differences influenced by pre-existing MBC and Abs.

Collectively, challenge induced serum Ab changes mostly paralleled B cell changes: CbΔ4 rHA induced Abs showed a greater “primary addiction” than SbΔ4 HA Abs, despite nearly identical antigenic distance from *wt* HA ^29^. We hypothesized that pre-existing Cb and Sb specific antibodies have distinct effects on secondary Ab responses to rHA immunization.

### Transferred monoclonal and polyclonal antibodies do not inhibit naïve B cell responses following vaccination with HA protein

The effects of pre-existing antibodies on B cell fates have been explored by mAb transfer ^26, 27, 28, 53, 54, 55^ in humans and animals. We examined the effect of mAbs targeting Cb (mAb H9-D3, green) or Sb (mAb H28-E23, yellow) on primary responses to rHA (Figure 4A)^35^. To equalize the effective dose, we transferred a standardized *wt* HA-binding Ab dose based on their ELISA maximum binding capacity (Bmax) (Supplementary Figure 5A), using biotinylated mAbs to enable their removal from sera in subsequent analysis of their effects on antibody responses to vaccination. We transferred 0.1xBmax (between 0.002 and 0.005 ug per dose), 1xBmax (between 16 and 18 ug per dose) of H9D3 and H28E23 intraperitoneally (i.p.) and 100µg (5.6x Bmax) of H28E23 (Figure 4A; Supplementary Figure 5A). Both 1x Bmax and 100µg doses are in the range of previous studies, with 100µg being the highest amount used by others and employed here as a positive control ^26, 27, 28^. Four hours post transfer, we vaccinated mice in the left footpad with *wt* rHA. Mice were sacrificed 14 days after vaccination to enable a proper GC response to develop (Figure 4A).

**Figure 4:**
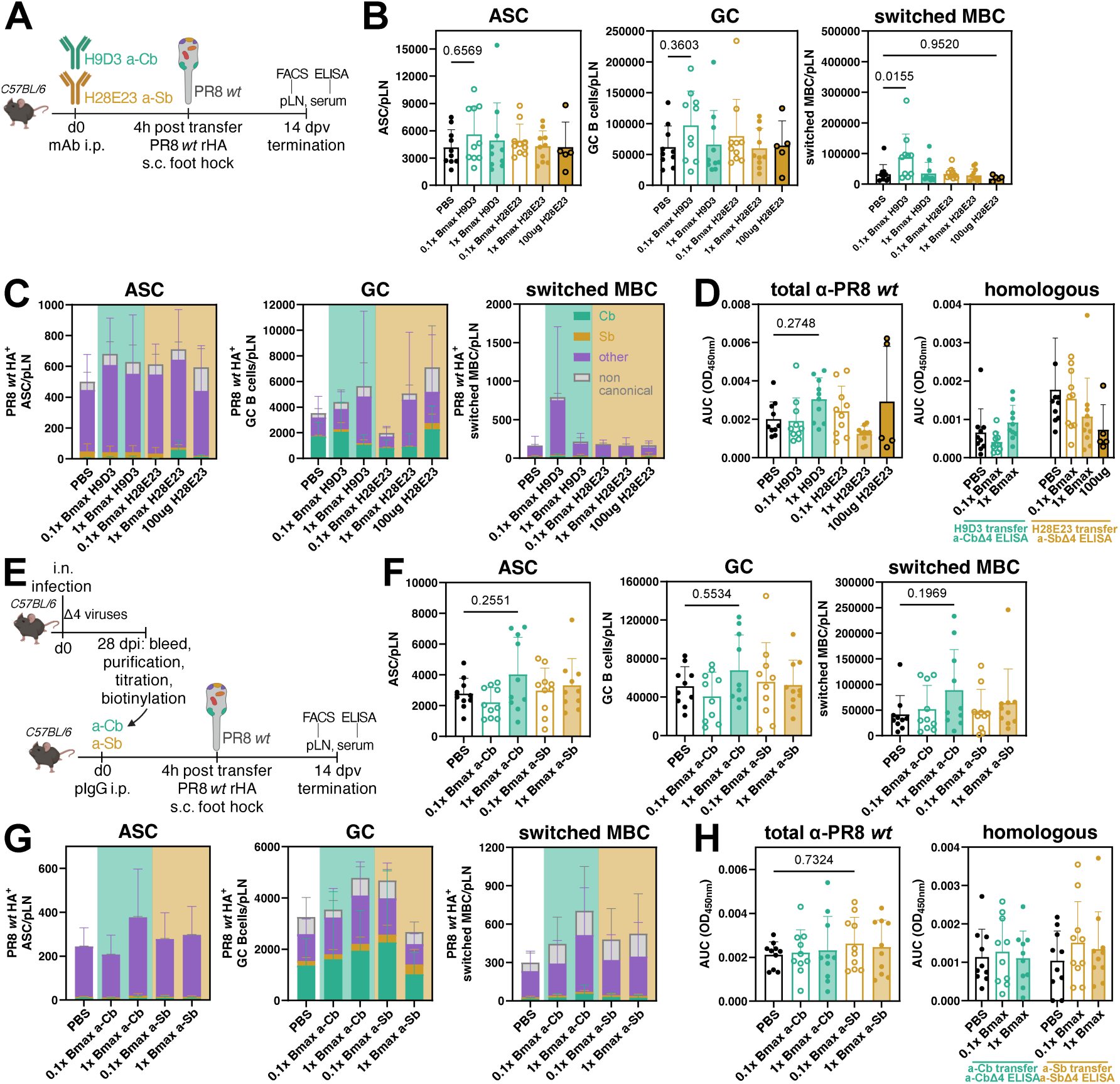
Transfer of mAbs and pIgG does not affect B cell responses to rHA vaccination. **A:** Experimental workflow for mAb transfer with rHA challenge are shown. **B:** Total cells per pLN at 14 days post vaccination (dpv) in ASC, GC B cells and switched MBC after mAb transfer. **C:** Total *wt* HA^+^ ASC, GC B cells, switched MBC and immunodominance patterns at 14dpv. The mAb transferred is indicated in color of plot background (green = a-Cb/H9D3; yellow = a-Sb/H28E23) **D:** AUC OD450nm values of titrated sera at 14dpv are shown, with *wt* HA used for coating (left), CbΔ4 and SbΔ4 used for coating (right) as mAb homologous antigens (CbΔ4 as H9D3 homologous antigen, SbΔ4 as H28E23 homologous antigen). **E:** Experimental workflow of pIgG transfer with rHA challenge are shown. **F:** Total cells per pLN at 14dpv in ASC, GC B cells and switched MBC after pIgG transfer with rHA challenge. **G:** Total *wt* HA^+^ ASC, GC B cells, switched MBC and immunodominance patterns at 14dpv. pIgG transferred is indicated in color of plot background (green = a-Cb; yellow = a-Sb). **H:** AUC OD450nm of sera at 14dpv are shown, with *wt* HA used for coating (left), CbΔ4 and SbΔ4 used for coating (right) as pIgG homologous antigens (CbΔ4 as a-Cb homologous antigen, SbΔ4 as a-Sb homologous antigen).p-values were calculated by ordinary one-way ANOVA, with all groups compared to PBS/mock transfer (**B, D, F, H**). (∗p < 0.05; ∗∗p < 0.01; ∗∗∗∗p < 0.0001). Lowest p-value indicated if no significance was detected. At least 2 experiments per group were performed. Figures represent data from two experiments with 3-5 mice per transfer group per experiment. All data are shown as mean ± SD.

We enumerated total and antigen-site specific ASC, GC and switched MBC in the draining pLN (Figure 4B-C; Supplementary Figure 5B). Surprisingly, despite using up to saturating doses of high-avidity mAbs, these had no significant effects on B-cell responses compared to PBS transfer, including total and antigen-specific GC B cells and changes in the antigenic site- specific immunodominance profile (Figure 4B-C). ELISA analysis extended *status quo* maintenance to serum Ab titers (Figure 4D).

We hypothesized that polyclonal Abs may be more effective in modulating B cell responses *via* Ab feedback ^24, 30, 52, 56, 57^. We purified polyclonal IgG (pIgG) from mouse serum 28 d post-i.n. PR8 or Δ4 virus infection using Melon Gel, which retains most non-IgG serum proteins, resulting in ∼ 90% purification of IgG, mainly IgG2b and IgG2c isotypes and lacking IgM (Figure 4E, Supplementary Figure 5C). To distinguish transferred from host Abs, we biotinylated pIgG before transfer (Figure 4E) and used rHA ELISA to determine the Bmax and titer of all anti-HA pIgG pools generated (Supplementary Figure 5D). Four hours post-i.p. transfer we detected biotinylated pIgG in sera by its binding to goat anti-mouse IgG in ELISA (Supplementary Figure 5E).

We then repeated the mAb (Figure 4E) experiment with CbΔ4- and SbΔ4-speific pIgG with near identical results. (Figure 4F-G), even when using pIgG to *wt* HA at high concentration (8x Bmax; 100µg) (Supplementary Figure 5F). Host serum anti-HA Ab binding remained virtually unchanged in transferred *vs* non transferred mice (Figure 4H). Transferred pIgG alone did not induced an anti-HA immune response, ruling out anti-Ig response (Supplementary Figure 5G). Using pIgG purified by protein G gave similar results (Supplementary Figure 5H).

Our data show that mAb and polyclonal Ab, at the concentrations used, do not affect B cell and Ab responses after vaccination with *wt* rHA, implicating that rather MBCs influenced the B cell and Ab differences we detailed in Figures 1 and 2.

### Immunogen valency dictates Ab modulation of naive B cell responses

The Ab transfer results are nevertheless surprising as numerous laboratories have reported that transferred mAbs or sera interfere with antigen specific *de novo* B cell responses ^24, 26, 27, 28, 52, 55, 58^. We noted that in most studies the challenging immunogen expressed multiple antigen copies on the surface of cells, virions, or artificial particles. rHA, as a monomeric (though homotrimeric) vaccine antigen, cannot easily crosslink the BCR on B cells ^59^. Furthermore, immune complex (IC) formation is disfavored using homotrimeric rHA, as a single Ab can most likely only bind one epitope per antigenic particle ^60, 61, 62, 63^. Focusing on GC and serum Ab responses (most studied previously) we examined the influence of antigen valency (and thus, indirectly, avidity) after passive Ab transfer, using three vaccine antigens of increasing valency (Figure 5A), in addition to rHA: 1) Streptavidin (SA) tetramerized biotinylated HA; 2) Split virion vaccines (following the protocol used for human influenza vaccines) consisting of HA present in a mixture of monovalent and multivalent HA aggregated by their hydrophobic tails, with up to 50 HA trimers per aggregate ^64^; and, 3) UV-inactivated virions, which have 500 or more HA trimers per virion ^65^.

**Figure 5:**
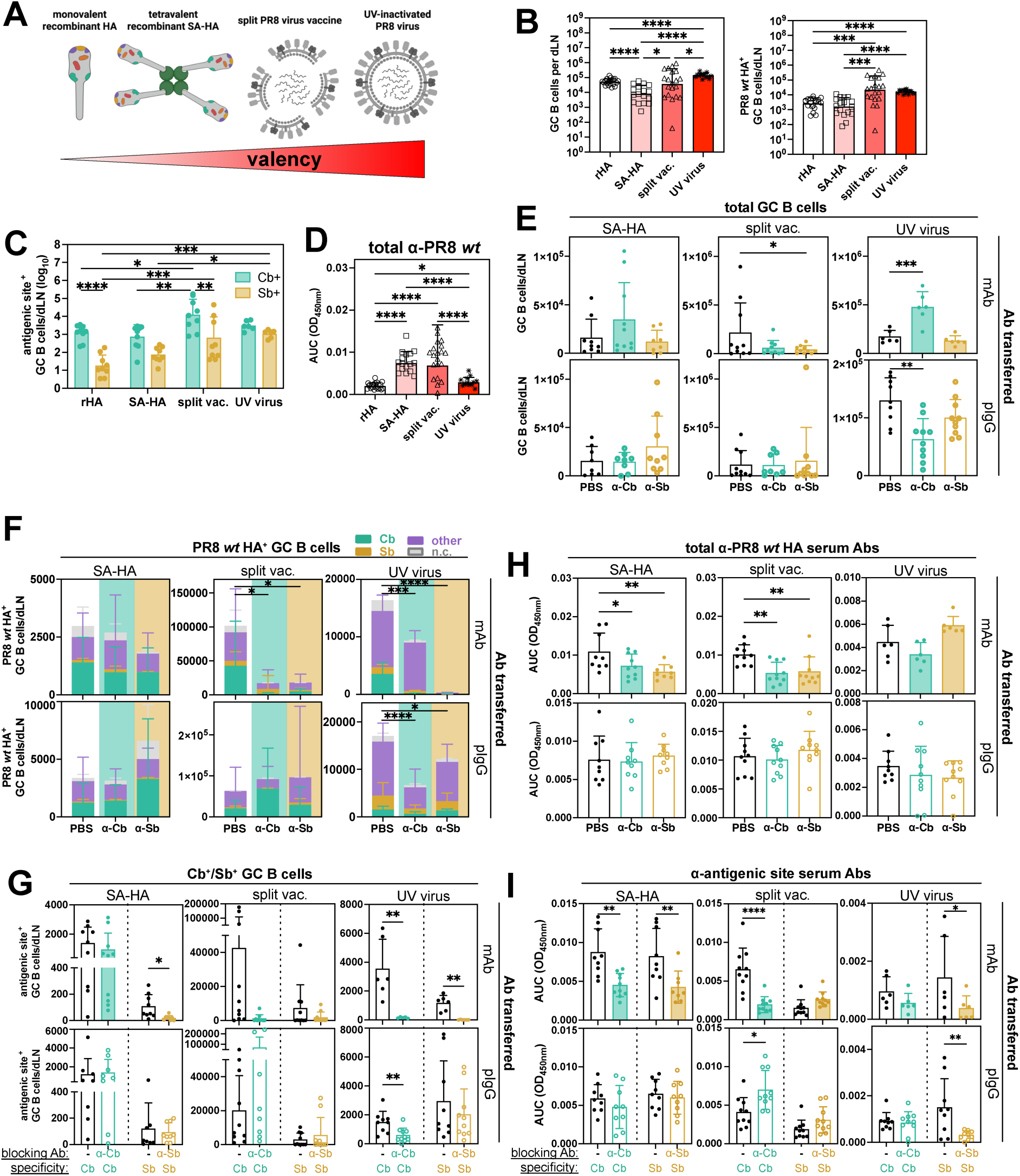
Ab modulation of B cell responses depends on antigen valency. **A:** Scheme displaying vaccines used in this work. **B:** Total GC B cells and PR8 *wt* HA^+^ GC B cells per dLN at 14 days post vaccination (dpv) after PBS (mock) transfer are shown. **C:** Total Cb^+^ and Sb^+^ GC B cells per dLN at 14dpv after PBS (mock) transfer are shown. **D:** AUC OD450nm determination of anti-*wt* HA^+^ IgG in serum at 14dpv after PBS (mock) transfer. Data for rHA vaccination is the same already reported in Figure 4 and shown again here for comparison. **E:** Total GC B cells per dLN at 14 dpv with or without mAb or pIgG transfer and for different vaccines are shown. **F:** Total *wt* HA^+^ GC B cells and immunodominance patterns at 14 days post vaccination with or without mAb or pIgG transfer and for different vaccines are shown. The mAb/pIgG transferred is indicated by color of plot background (green = a-Cb; yellow = a- Sb). **G:** Cb^+^ and Sb^+^ HA^+^ GC B cells at 14 dpv with or without mAb or pIgG transfer and for different vaccines are shown. **H:** AUC OD450nm of sera at 14 days post vaccination with or without mAb or pIgG transfer and for different vaccines. *wt* HA was used for coating. **I:** Calculated AUC OD450nm of sera at 14 dpv with or without mAb or pIgG transfer and for different vaccines are shown. CbΔ4 was used for coating to define Cb-specificity while SbΔ4 was used for coating to define Sb-specificity. p-values were determined by one-way ANOVA (**B, D, E, F, F**), unpaired t-test (**G, I**) or two-way ANOVA (**C**). (∗p < 0.05; ∗∗p < 0.01; ∗∗∗p < 0.001; ∗∗∗∗p < 0.0001). Figures represent data from two experiments with 3-5 mice per transfer group per experiment. All data as mean ± SD. The dose used for transfer was 1x Bmax for mAb and 0.1x Bmax for pIgG.

Two weeks post vaccination, in animals with no passively transferred Abs, split or inactivated virus vaccines (but not SA-HA) increased GC size and numbers of *wt* HA^+^ specific GC B cells in dLN, relative to rHA vaccination (Figure 5B). This could be explained by increased valency but also by the extra adjuvanticity from residual viral components. Virion-based vaccines were better at inducing Cb- and Sb- binding GC B cells while generally retaining Cb dominance (Figure 5C). Similarly, all multivalent vaccine antigens were better than monovalent rHA at inducing serum anti-HA Ab responses (Figure 5D). Surprisingly, HA-titers were comparatively lower when vaccinating with UV-inactivated whole virus, possibly due to competition from NP and chicken host component (Figure 5D).

Having established baseline responses to the four vaccines, we transferred mAb (1x Bmax) or pIgG (0.1x Bmax) specific for either Cb or Sb sites, vaccinated mice with each of the vaccines, and analyzed B cell immunity two weeks post-immunization. Overall, transferred Abs had limited effects on the total magnitude of GC responses with few exceptions (Figure 5E). However, *wt* HA^+^ GC B cell responses induced by UV-virus vaccine were significantly decreased by all Ab treatments (Figure 5F). The *wt* HA^+^ GC B cell responses after split influenza vaccine were also inhibited by mAbs, but not pIgG, possibly due to the lower avidity (determined by resistance to elution by 6M urea in ELISA) or dose of polyclonal IgG (Figure 5F and Supplementary Figure 5I). Of note, Cb-specific pIgG had a higher avidity index compared to Sb pIgG (Supplementary Figure 5I). Effect on GC B cell immunodominance was observed within the same groups, with passively transferred Abs potently blocking responses to the cognate antigenic site (Figure 5F-G). The cognate inhibition was even observed for Sb-specific responses after Sb-mAb administration and SA-HA vaccine (Figure 5G). Total serum Abs to *wt* HA were generally only affected by mAb but not pIgG administration (Figure 5H). Similarly, we observed a more potent antigenic-site specific suppression when using mAbs, independently of the vaccine (Figure 5I).

Taken together, these finding indicate that antigen valency is a major determinant of Ab-mediated inhibition of B cell and Ab responses. While mAbs and pIgG had little effects on rHA vaccination, they blocked overall and antigenic site-specific B cells when vaccine valency was increased.

### MBC suppress germinal center recruitment of antigen-specific naïve B cell

The experiments above demonstrated that while previous antigen exposure influenced recall responses markedly, Abs alone were unable to reshape humoral immunity to low valency antigens. In a more physiological scenario, memory responses will take place in the presence of both Abs and MBC, and the antigen would be multivalent. We hypothesized that since MBC can rapidly differentiate into ASC secreting Abs in situ ^13, 66^, they may be able to limit naïve B cell recruitment into dLN GCs more efficiently than circulating Abs, due to high local concentrations. Consistent with this idea, Abs were readily detected in pLN of i.n. infected, footpad vaccinated mice, 7 days after boosting (Figure 6A).

**Figure 6:**
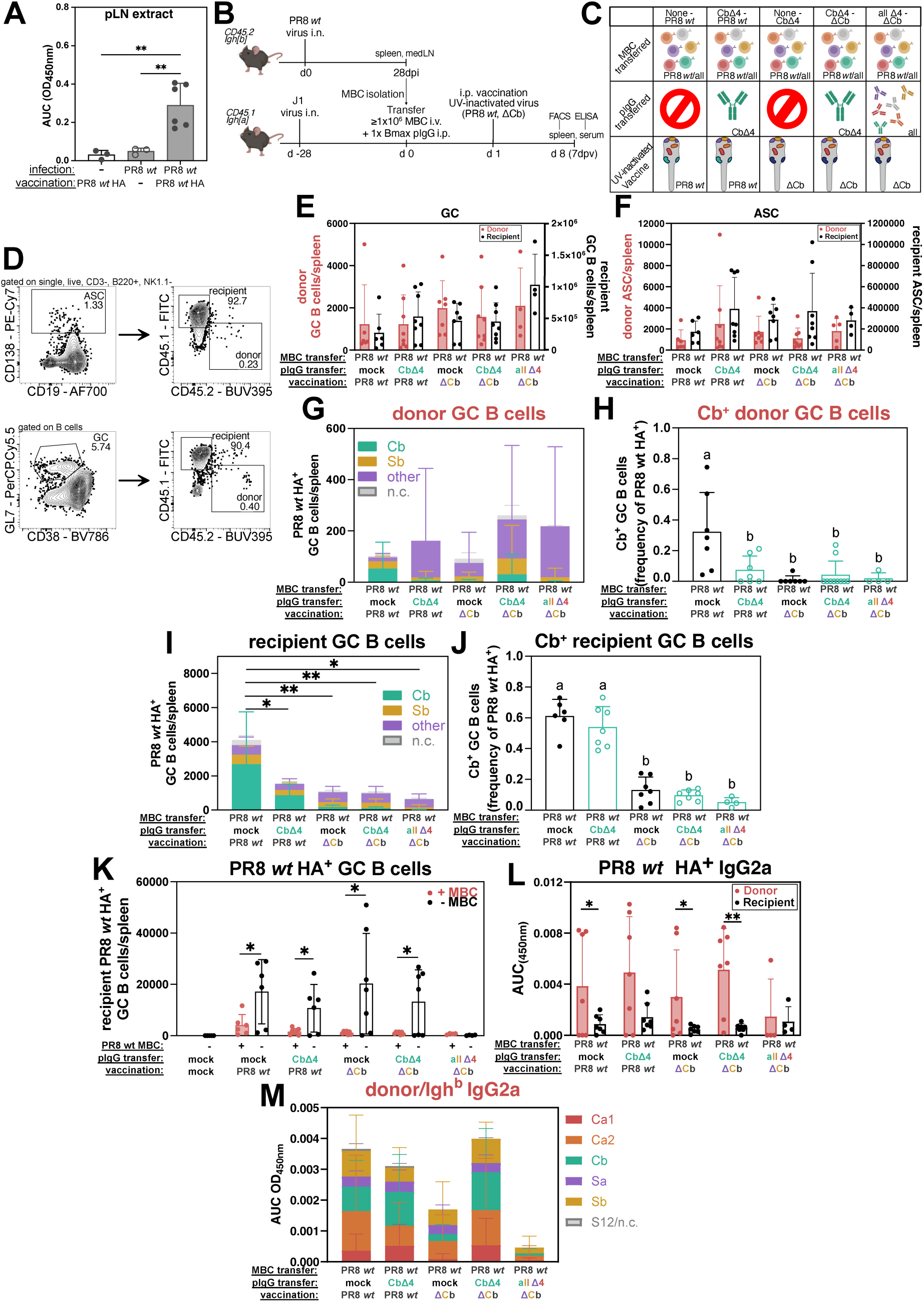
Differential effect of antibodies on naïve B cells and MBC fate and immunodominance and MBC-mediated suppression of antigen-specific naïve B cell. **A:** Local Ab at 7 days post vaccination (dpc) in the pLN as measured by ELISA. **B:** Experimental scheme of MBC + pIgG adoptive transfer into recipients are shown. **C:** Groups of pIgG and vaccination combinations used are demonstrated. **D:** Representative FACS plots of single, live, CD3^-^ NK1.1^-^ B220^+^ cells showing identification of donor and recipient cells among ASC (top) and GC B cells (bottom). **E:** Total count of recipient *wt* HA^+^ GC B cells in spleen with and without MBC transfer are shown. **F:** Total count of donor and recipient GC B cells in spleen and **G:** ASC in spleen. **H:** Immunodominance patterns of donor *wt* HA^+^ GC B cells and **I:** frequency of Cb-specific GC B cells, among *wt* HA^+^ cells in spleen at 7dpc**. J:** Immunodominance patterns of recipient *wt* HA^+^ GC B cells and **K:** frequency of Cb-specific GC B cells, among *wt* HA^+^ cells in spleen at 7 days post vaccination. **L:** AUC OD450nm of recipient (Igh[b]) and donor (Igh[a]) *wt* HA specific IgG2a. **M:** Immunodominance patterns among *wt* HA binding IgG2a of donor antibodies. p-values were calculated by one-way ANOVA (**A, I, J, K**) or unpaired t test (**E, L**). (∗p < 0.05; ∗∗p < 0.01; ∗∗∗∗p < 0.0001). For **I** and **K** statistical differences are indicated by compact letter display (cld), where groups with different letters are statistically different. Figures represent data from n=2 experiments/group with 3-7 mice per challenge group per experiment. All data shown as mean ± SD. Animals with ≤50 donor *wt* HA+ B cells (ASC+IgM MBC+switched MBC+ GC B cells) per spleen were excluded from the analysis.

To address the relative contribution of MBC and pIgG to the results observed in the rechallenge experiment (Figures 1-3), we set up a system where both Abs and MBC were transferred into recipient mice. We used *C57BL/6* mice, expressing the congenic marker CD45.2 on immune cells and secreting Abs of the Igh[b] allotype, as donors and *B6 CD45.1 x Igh^a^* mice, expressing the congenic marker CD45.1 on cells and secreting Abs of the Igh[a] allotype, as recipients (Figure 6B). Total MBC that had specificities to all antigenic sites were purified from mice i.n. infected with *wt* PR8 before transfer (Figure 6B-C; Supplementary Figure 6A). For transfer of pIgG we either used pIgG purified from mice previously infected with CbΔ4 virus (thus only carrying Ab against the the Cb site of *wt* PR8 HA) or an equimolar amount of pooled pIgG from mice that had been individually infected with all 5 viruses carrying different Δ4 mutations (thus targeting all *wt* PR8 HA sites) (Figure 6C). As vaccines we used either *wt* PR8 or ΔCb (carrying all canonical antigenic sites, except for Cb).

We focused the experiments on UV virus vaccine, as this was the most sensitive to inhibition by mAb and pIgG transfers in the previous experiments (Figure 5F-G), and on the response to the Cb site because of its immunodominance ^29^. When setting up the experiments, we noted that recipient mice had to be pre-infected i.n. with J1, a reassortant PR8 virus expressing H3 HA (Supplementary Figure 6B-D) to enable transferred MBC to survive. J1 infected mice did not develop detectable Abs reactive against *wt* PR8 HA that could influence MBC responses against this antigen (Supplementary Figure 6D). Pre-infection was possibly needed to generate CD4 T cells targeting conserved internal viral antigens^67, 68^ or cross-reactive HA peptides^69^. Whatever the case, as J1 priming was an essential step to support transferred MBC survival and involvement in the recall response all experiments were performed in primed recipient mice (see Methods and Supplementary Figure 6B-D). Four weeks later, MBC and/or pIgG were transferred and a day later mice were vaccinated i.p with UV-inactivated virus (Figure 6B). Seven days post vaccination, both donor and recipient GC and ASC could be identified in spleen and medLN (Figure 6D and Supplementary Figure 6E). In this setup, engraftment was observed in the majority of transferred mice and detection of transferred B cells was strictly dependent on vaccination, demonstrating antigen-dependent expansion of MBC after immunization (Supplementary Figure 7A).

First, we determined the effect of MBC transfer alone on immune response. When compared to no transfer, the MBC transfer resulted in a general increase in overall GC and ASC cellularity (Supplementary Figure 7B-C). However, while the total number of activated cells increased, the number of *wt* HA^+^ naïve-derived GC and ASC cells decreased drastically in MBC- transferred *vs* non-transferred mice (Figure 6E and Supplementary Figure 7D-E), demonstrating that MBC alone could inhibit antigen-specific naïve B cell differentiation. We hypothesize that this observation was due to high levels of Ab being secreted by differentiated donor MBC *in situ* (Figure 6A).

Subsequently, we analyzed the humoral response of mice receiving both MBC and pIgG. Groups were designed so that MBC repertoire was broad and Ab response narrow ^44^. Overall, recipient cells dominated the response, but transferred cells were also detected (Figure 6F-G and Supplementary Figure 7F-G). Consistent with our other data above (Figure 5), the total numbers of responding GC and ASC cells were not influenced by co-transfer of Abs (Figure 6F-G and Supplementary Figure 7F-G). However, when we defined antigen specificity as in previous experiments (Figure 2A), we observed subtle differences in *wt* HA specific donor MBC that could re-enter the GC reaction, depending on the presence of transferred pIgG (Figure 6H). Specifically, we found that pIgG blocked donor B cell response to the homologous antigenic site they targeted (e.g. Cb-specific Abs decreased the MBC-derived response to Cb after PR8 *wt* virus infection) (Figure 6H-I). Interestingly, this was not the case for *de novo* responding B cells, as recipient (naïve-derived) *wt* HA^+^ GC B cells were largely Cb-site specific to the same extent even after CbΔ4 pIgG transfer, albeit the total response was lower when compared to no pIgG-transfer (Figure 6J-K). Similarly, Cb-specific, donor (MBC- derived) ASC were more efficiently inhibited by Cb-reactive pIgG when compared to recipient (naïve-derived) (Supplementary Figure 7H-K). However, as noted above, the presence of MBC alone also reshaped the immunodominance patterns of naïve-derived B cells (Supplementary Figure 7D-E). The overall results suggest differential Ab feedback and MBC regulation of naïve B cells *vs* MBC.

We finally measured Abs secreted by recipient (Igh[a]) vs donor (Igh[b]) ASC (Figure 6L-M). At this early time point (day 7 post vaccination), donor Ab titers (Igh[b]) were higher than the recipient Ab titers (Igh[a]) (Figure 6L), showing that early secondary Ab responses from ASC are dominated by activation of antigen experienced MBC. When analyzing immunodominance in donor cells the pattern was similar across conditions, consistent with a general reactivation of all MBC, with few exceptions (Figure 6M). For example, when ΔCb vaccine was used without Ab presence, there was very limited production of Cb-specific Abs, as expected; and a combination of all pIgG blocked differentiation of almost all MBC to ASC.

Based on these findings, we conclude that pre-existing Abs impact MBC and naïve cells in different ways and that MBC themselves also affect the magnitude of total and antigen specific *de novo* B cell responses, likely through rapid differentiation and local Ab secretion.

## DISCUSSION

Using cell fate tracking experiments and transfer of epitope specific Abs and MBC, we revealed how each component influences MBC and naïve B cell responses to vaccination with drifted antigens in naïve and infection-experienced animals. We have made 4 key discoveries: 1) Even a few amino acids substitutions in immunodominant vaccine antigenic sites, comparable to normal antigenic drift, can reshape naïve B cell recruitment into GC and secondary B cell responses, 2) Ab-mediated regulation of *de novo* B cell responses depends on antigen multivalency, 3) MBC can suppress naïve B cell entry into GC in an antigen-specific manner, and 4) pre-existing Abs differently regulate MBC and naïve B cells.

Schiepers *et al.* used drifted monovalent rHAs representing various IAV strains to examine secondary MBC responses with protein priming and boosting in opposite footpad. They found that boosting with drifted, but not identical, rHA resulted in increased numbers of HA-binding GC B cells and increased frequency of MBCs re-entering into secondary GCs in dLNs ^23^. The boosting rHAs were, however, extremely drifted from priming rHA, carrying 10%-20% amino-acid substitutions, which represents decades of normal antigenic drift in influenza. To create a closer to a real-life human scenario we here used engineered Δ4 PR8 rHAs to mimic the typical circumstance of being challenged with an IAV strain (by infection or vaccination) that has only drifted a few years from viral strains used for prior infection. Despite this and our use of i.n. infection for priming, we also observed that secondary GCs are mostly populated by *de novo* responses, with the proportion of GC cells derived from MBC re-entry increasing up to ∼20% if only considering HA-specific cells. However, we may be underestimating the memory contribution, as early GC-independent, and thus non-fate mapped MBC, may also participate in recall responses ^10^. We also found that most fate-mapped MBC rapidly differentiated into ASC in the dLN after vaccination. Even so, we acknowledge that the number of HA-specific ASC likely were even higher, as terminally differentiated IgG producing PC mostly lack surface BCR ^70^. Importantly, the degree of drift in boosting vaccine rHA governed the magnitude of both the MBC response to the boosted site but also the naïve B cell responses to the novel antigenic sites, resulting from drift. This dependency on the degree and nature of antigenic drift likely explain some of the contradictory results in prior original antigenic sin studies ^47^.

Several recent studies examined the effects of mAbs, pIgG and whole sera on *de novo* activation of B cells specific for the same epitope targeted by the Ab upon vaccination ^24, 26, 27, 28, 30, 52, 53, 54, 56, 57, 71^. The results from these studies should be reevaluated in light of our finding that Ab feedback inhibition depends not only on the nature of the antigen but also on its valency. GC responses to rHA vaccination were mostly unperturbed by transferred Abs but likely affected by MBC, which we expect to act by intranodal Ab secretion after rapid differentiation to ASC. This implies that transferred Abs possibly do not achieve sufficient concentration in B cell follicles to affect B cell activation, unless multivalent vaccine antigens are used, likely due to the increase in functional Ab avidity in this case. Furthermore, ICs preferentially accumulate on FDCs when formed with multivalent antigens ^72^, thereby allowing for sustained antigen availability within GC to amplify B cell responses (Figure 5B). This provides a possible mechanistic explanation to our findings: low-valency antigens do not form sufficiently stable ICs to deposit on FDCs, and circulating antibodies remain too diluted in the follicle. In contrast, multivalent vaccines generate high-avidity ICs that can saturate epitopes, concentrate on FDCs, and effectively mask antigens from competing B cells.

Lack of Ab inhibition of monovalent vaccination echoes a recent study from Termote *et al.*^20^, but contrasts with Dvorscek *et al.* ^26^ who found that transferred mAbs could block responses to the monovalent protein hen egg lysozyme given that they had sufficient affinity. Differences between our studies is that while we examined responses from endogenous polyclonal B cells of varying specificity and avidity, Dvorscek *et al.* used identical mAb and fixed monoclonal B cells competing with each other for antigen access. Furthermore, the affinity of OVA-specific Abs ^26^ is overall higher than avidity of B cells and Abs to HA ^73^. The discrepancies highlight that this is a tightly regulated system, where modulation of antigen valency as well as affinity may help to overcome Ab-dependent inhibition. Finally, timing of Ab administration may also be crucial as immune serum (containing also IgM) transferred few days after immunization was also able to modulate immunity to monovalent antigens ^30, 52^.

Our experimental design enabled precise characterization of epitope-specific MBC activation and dissection of differences in specificity between naïve B cells vs reactivated MBC as well as their secreted Abs. While Schiepers “primary addiction” model ^22^ is consistent with some of our findings, we find that monovalent rHA vaccines, even if differing from priming HA by only 5 amino acids, can profoundly reshape secondary B cell responses (Figure 1-3 and 6). Importantly, our MBC transfer experiments (Figure 6E) establish that that MBC play a critical role in B cell fate and the resulting immunodominance of booster vaccine even for *de novo* responses initiated after vaccination. This effect can likely be explained by rapid differentiation of MBC to ASC with the local secretion of enormous amounts of blocking Abs (5 – 10×10^7^ molecules per cell per hour ^74, 75^) *in situ*. Interestingly, our model is consistent with results from Schiepers *et al.* where, by deleting *Prdm1* in GC-derived cells (including MBC), they found decreased Ab-mediated suppression ^23^.

Our transfer studies highlight the complex interplay between polyclonal Abs and B cells. Using MBC and Abs with a range of avidities and specificities we recreated complex scenarios of pre-immunity similar to what occurs with IAV, CoVs and other drifting human viruses ^43, 44, 45^. In this context, we found that, using a multivalent antigen vaccine, epitope-specific MBC are more likely to re-enter a GC in the absence of pre-existing Abs (Figure 6H) ^23^. This Ab-mediated inhibition was more potent in blocking GC entry of MBC as compared to naïve B cells, thus explaining results from the infection/vaccination experiment (Figure 1) and from previous studies ^21, 22, 23^. These scenarios unveiled how pre-existing MBC, Abs and challenge antigen interact to determine MBC and naïve cells activation and magnitude as well as antigenic site specificity of the overall response.

A limitation of our study is that, while it considered multiple factors simultaneously and allowed a more faithful recapitulation of real life scenarios ^43, 44, 45^, by design, it could not resolve the detailed affinity/avidity and epitope-specificity of Abs *vs* B cells, as was possible in some previous studies ^22, 23, 24, 26, 27, 28^. Furthermore, to capture reactivation of MBC we conducted our experiments at 7 days post vaccination. Future studies need to be conducted to gain further insight into earlier and later timepoints to illustrate how secondary B cell responses are initiated and persist.

In summary, our findings illuminate the complex regulation of secondary B cell responses and original antigenic sin phenomenon and should be taken into consideration when designing monovalent and multivalent immunogens for vaccination.

## ACKNOWLEDGMENTS

We thank all members of the Angeletti lab for helpful discussion and helpful interactions. Additionally, we want to thank the staff or the animal facility Experimental Biomedicine, especially Pernilla Ahlgren, for excellent upkeeping and care of animals used in this study. We also want to thank the Protein Production Sweden (Gothenburg, Sweden), especially Malin Bäckström and Mikael Andersson, for production of rHA proteins. We thank Tomohiro Kurosaki (WPI Immunology Frontier Research Center, University of Osaka, Japan) for kindly providing *S1pr2-ERT2-cre-TdTomato* mice ^51^ and Masaru Kanekiyo (VRC, NIAD, NIH) for the plasmids for recombinant S12 HA expression.

The study was supported by grants from the European Research Council (ERC-StG, B-DOMINANCE, grant no. 850638 to DA); the Swedish Research Council (grant no. 2017- 01439, 2021-01164, 2021-01165 to DA); the Knut and Alice Wallenberg Foundation (grant no 2021.0033 to DA), the Jeanssons Foundation (grant nos. JS2018-0011 and JS2019-0038), the Claes Groschinsky Foundation (grant no. M18237). NRM is supported by Svenska Sällskapet för Medicinsk Forskning post-doctoral grant (grant no: PD20-0017). BioRender was used to generate images and graphics.

## AUTHOR CONTRIBUTIONS

L.R. Performed the majority of experimental work; K.S. produced virus, handled animal breeding, performed TCID_50_ assays as well as some vaccinations and ELISA; D.A. performed flow cytometric comparison of multivalent vaccines; N.R.M. started optimization work for MBC transfer; D.F.B. contributed to protein production; D.A., I.K., J.S.G., M.C.M were involved in development of new HA escape mutants; J.W.Y. provided key reagents; D.A. conceptualized the study; L.R., D.A. and M.B. designed all experiments and interpreted data; L.R. and D.A. wrote the first draft of the manuscript with critical input from J.W.Y. and M.B.. All authors reviewed, edited and approved the manuscript.

## MATERIALS AND METHODS

### Animals

Wild type C57BL/6 mice were obtained from Janvier Labs. *S1pr2^CreERT^*^2^ animals were kindly provided by Tomohiro Kurosaki ^51^ and crossed with *Ai14 R26^Tomato^* to obtain *S1pr2^CreERT^*^2^*- R26^tdTomato^*. Recipients for MBC transfer, PepBoyIgha, were bred onsite from *B6.CgGpi1aThy1a Igha/J* and *B6.SJLPtprcaPepcb/BoyJ* animals obtained from Jackson Laboratory. Animals were kept in a specific pathogen free facility under Biosafety-Level 2 conditions. Breeding was conducted under the ethical permit 3307/20 and experiments under the ethical permits 1666/19, 2230/21 and 38/23. All ethical permits were approved by the Swedish Board of Agriculture.

For cell fate tracking experiments*, S1pr2-ERT2-cre-TdTomato* mice were infected with 25 TCID50 PR8. Every second day, mice received an oral gavage of 2mg tamoxifen per 100ul of corn oil. At 26 days post infection (dpi), treatment was stopped, at 27dpi animals were bled from vena saphena and at 28dpi animals were challenged as indicated.

7 days post vaccination (dpv), animals were sacrificed. Mediastinal lymph node (medLN), popliteal lymph node (pLN) and blood for serum were harvested. Tissue was processed for flow cytometric staining and serum used for ELISA.

### Viruses and proteins

Viruses were grown in 10 days old embryonated chicken eggs or propagated in MDCK cell culture and purified using ultracentrifugation on a sucrose gradient. Viruses used were A/PR/8/34 (PR8), its Δ4 and Δ1 escape mutants. For the new Δ4 selection, we reselected the original Δ4 viruses using a mixture of 2 mAbs targeting the Sb cryptic epitopes defined by residues 189 and 198 (H1 numbering)^50^, according to the same protocol previously described ^29^. Resulting viruses had the same mutations as described ^29^ but also carried the following extra mutations: SaΔ4 K189E, Ca1Δ4 E198G, Ca2Δ4 E198G and CbΔ4 Y201S. Escape was confirmed by ELISA. ΔCb was a precursor of Δ4 viruses and defined by the following mutations: L75P, V77M, R78K, E124G. J1, is a PR8 reassortant virus expressing an H3N1 instead of the PR8 HA. Viral titers were determined by TCID_50_.

Virus was UV inactivated on ice for 20-30 min and stored at -20°C. HAU was determined from UV inactivated virus by hemagglutination inhibition (HI) assay as described previously ^29^. Briefly, a 1% solution of chicken red blood cells was incubated in a 96-well round bottom plate with a serial titration of influenza A virus for 30min at RT. HAU titer was determined subsequently.

Recombinant HA (rHA) proteins were produced by Protein Production Sweden as previously described ^76^. All rHA included the Y98F mutation to abrogate sialic-acid binding. Protein biotinylating was performed using BirA-mediated enzymatic reaction, according to manufactureŕs instructions (Avidity LLC).

### Infections and immunizations

For i.n. infection, animals were anesthetized and 25-100 TCID50 of virus per 25µl HBSS/0.1%BSA were injected in one nostril. Animals were monitored for signs of severe disease and sacrificed if too sick. Animals were kept for 28 days to allow sufficient B cell memory and Ab titers to develop. To check titers, animals were bled 26 days post vaccination from the Vena Saphena.

For rHA immunizations, 10µg of rHA was mixed with AddaVax™ (MF59®- like squalene oil-in-water adjuvant, InvivoGen) in a 1:1 v/v ratio to 30µl and injected subcutaneous (s.c.) into the left foot hock.

For SA-HA immunization: biotinylated HA was tetramerized with 4 fold molar excess of SA on ice and stored at 4 °C. 10µg of the complex was mixed with AddaVax™ (MF59®- like squalene oil-in-water adjuvant, InvivoGen) in a 1:1 v/v ratio to 30ul and injected s.c. into the left foot hock.

For split virus vaccine: purified virus was fractionated by incubation with an equal volume of 15% octyl-β-glucoside. After addition of PBS, the solution was spun at 50,000g for 2 h at 4 °C. The supernatant, containing HA was collected and quantified using a Bradford assay. HA presence and amount was confirmed using immunoblot with HA_2_-specific mAb. 8µg of split vaccine was mixed with AddaVax™ (MF59®- like squalene oil-in-water adjuvant, InvivoGen) in a 1:1 v/v ratio to 30ul and injected s.c. into the left foot hock.

For UV inactivated virus immunizations, 2000 HAU were injected either intraperitoneal (i.p.) 100ul in PBS or s.c. in the foot hock 30µl in PBS.

### ELISA with serum, purified Abs or LN supernatant

96 half-well plates for high protein binding were coated with rHA, recombinant Neuraminidase (rNA) from PR8, recombinant Nucleoprotein from PR8 (rNP) or UV-inactivated J1 over night (o/n) or up to 1 week. mAbs, Fabs and pIgG were diluted to a defined concentration and sera were diluted 1:100 in PBS-T. Diluted samples were titrated 2-fold down from the first row and incubated for 1.5h. After that, a secondary antibody coupled to peroxidase was used, incubating for 1h. Plates were finally developed, adding TMB for 5min and stopping the reaction with 2M H_2_SO_4_. Pates were read at 450nm within 30min of development.

For avidity ELISA, plates were coated with *wt* HA and pIgG was used in duplicate at 1xBmax. Following incubation, unbound pIgG was removed by treatment with either 6 M urea or PBS for 5 min to disrupt low-avidity interactions. Secondary antibody incubation and plate development were performed as described above. The avidity index was calculated as the ratio of absorbance in urea-treated wells to that in ÅBS-treated controls.

### Preparation of HA probes and staining of tissue samples for flow cytometry

rHA was biotinylated with the Biotin Protein Ligase standard reaction kit (Avidity), excess biotin was removed using a 30kDa MWCO protein concentrator. Biotinylated rHA was conjugated to streptavidin (SA) coupled to fluorescent molecules to create HA probes. Once conjugated, rHA was stored at 4°C in the dark.

Harvested tissue was either grinded on a 70µm filter or smashed directly in the tube. After washing with FACS buffer, samples were stained with fluorescent labelled mAbs in FACS buffer. Following a wash in PBS-EDTA, samples were stained for viability using LIVE/DEAD™ Fixable Aqua Dead Cell Stain Kit or LIVE/DEAD™ Fixable Far Red Dead Cell Stain Kit (Thermo Fisher). Subsequently, samples were fixed using 1.5% PFA in PBS and stored at 4°C in the dark until acquisition (maximum 1 week). Just before acquisition, 10µl of counting beads were added (SONY).

Samples were acquired on a BD LSR Fortessa X-20 (BD Biosciences) or ID7000 (SONY) and analysed using FlowJo V.10 software (BD Biosciences).

**Table 1:**
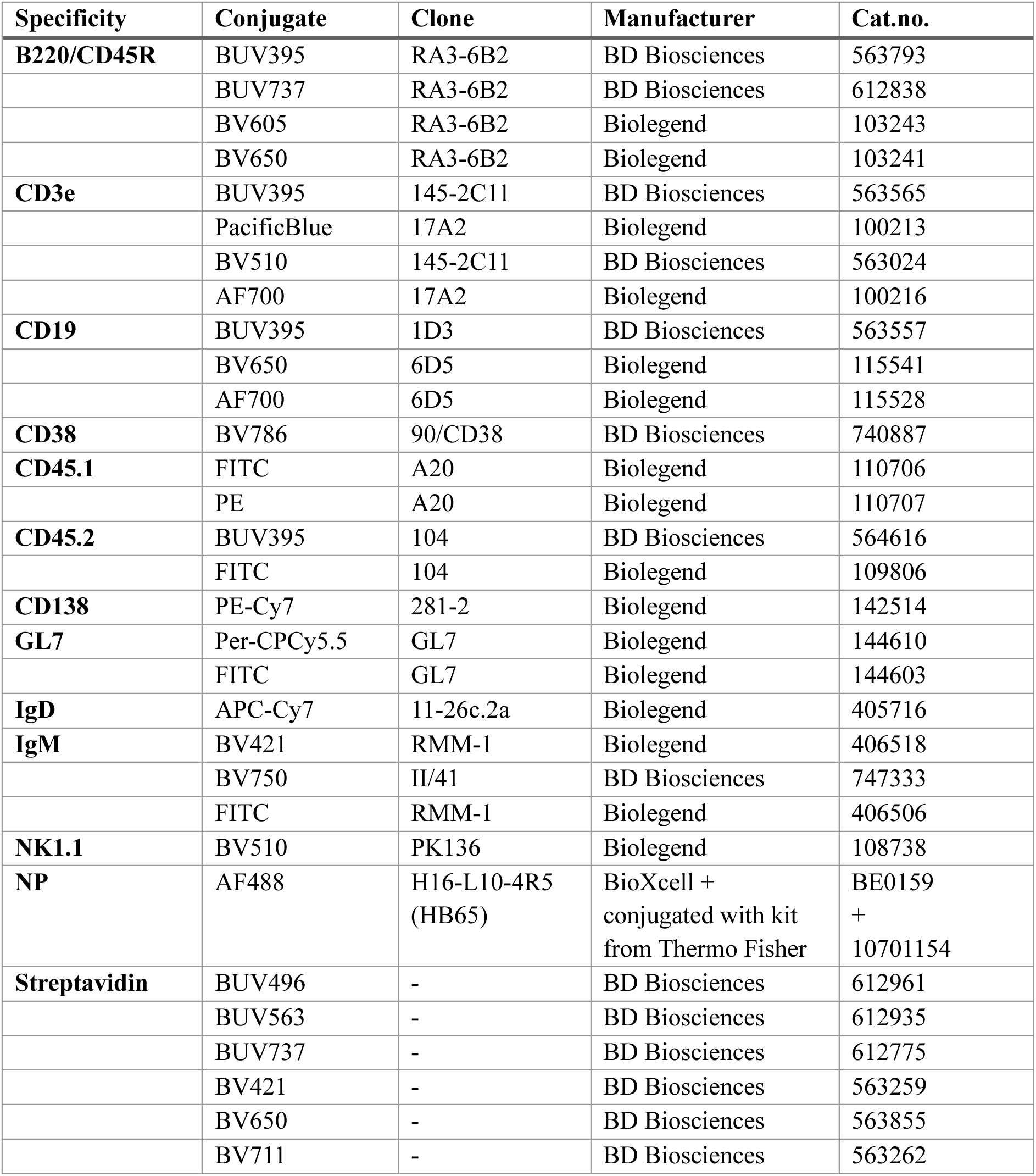
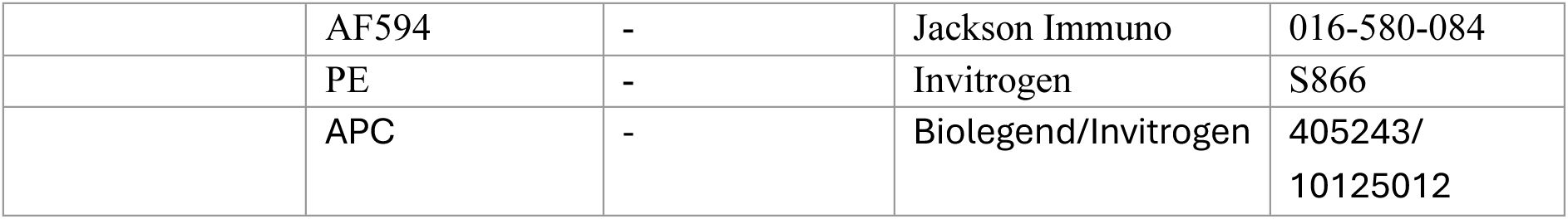
Fluorescent labelled antibodies used for flow cytometric staining.

| Specificity | Conjugate | Clone | Manufacturer | Cat.no. |
| --- | --- | --- | --- | --- |
| **B220/CD45R** | BUV395 | RA3-6B2 | BD Biosciences | 563793 |
|  | BUV737 | RA3-6B2 | BD Biosciences | 612838 |
|  | BV605 | RA3-6B2 | Biolegend | 103243 |
|  | BV650 | RA3-6B2 | Biolegend | 103241 |
| **CD3e** | BUV395 | 145-2C11 | BD Biosciences | 563565 |
|  | PacificBlue | 17A2 | Biolegend | 100213 |
|  | BV510 | 145-2C11 | BD Biosciences | 563024 |
|  | AF700 | 17A2 | Biolegend | 100216 |
| **CD19** | BUV395 | 1D3 | BD Biosciences | 563557 |
|  | BV650 | 6D5 | Biolegend | 115541 |
|  | AF700 | 6D5 | Biolegend | 115528 |
| **CD38** | BV786 | 90/CD38 | BD Biosciences | 740887 |
| **CD45.1** | FITC | A20 | Biolegend | 110706 |
|  | PE | A20 | Biolegend | 110707 |
| **CD45.2** | BUV395 | 104 | BD Biosciences | 564616 |
|  | FITC | 104 | Biolegend | 109806 |
| **CD138** | PE-Cy7 | 281-2 | Biolegend | 142514 |
| **GL7** | Per-CPCy5.5 | GL7 | Biolegend | 144610 |
|  | FITC | GL7 | Biolegend | 144603 |
| **IgD** | APC-Cy7 | 11-26c.2a | Biolegend | 405716 |
| **IgM** | BV421 | RMM-1 | Biolegend | 406518 |
|  | BV750 | II/41 | BD Biosciences | 747333 |
|  | FITC | RMM-1 | Biolegend | 406506 |
| **NK1.1** | BV510 | PK136 | Biolegend | 108738 |
| **NP** | AF488 | H16-L10-4R5 (HB65) | BioXcell + conjugated with kit from Thermo Fisher | BE0159 + 10701154 |
| **Streptavidin** | BUV496 | - | BD Biosciences | 612961 |
|  | BUV563 | - | BD Biosciences | 612935 |
|  | BUV737 | - | BD Biosciences | 612775 |
|  | BV421 | - | BD Biosciences | 563259 |
|  | BV650 | - | BD Biosciences | 563855 |
|  | BV711 | - | BD Biosciences | 563262 |
|  | AF594 | - | Jackson Immuno | 016-580-084 |
|  | PE | - | Invitrogen | S866 |
|  | APC | - | Biologend/Invitrogen | 405243/<br>10125012 |

### Preparation of Abs and transfer

To obtain polyclonal IgG (pIgG), female C57BL/6 mice were infected with 50 TCID50 of PR8 virus. At 28dpi, animals were sacrificed with a terminal bleed during ketamine anesthesia. Sera were pooled and IgG was purified using a Melon™ Gel (Thermo Fisher) column on the ÄKTA Start chromatography system (Cytvia) according to manufacturer’s instructions. Obtained IgG was concentrated using a 50kDa MWCO protein concentrator.

To be able to identify transferred Abs quickly and to be able to remove them from a mix with donor Abs, they were biotinylated. The EZ-Link™ Sulfo-NHS-LC-Biotin, No-Weigh™ Format kit (Thermo Fisher) was used according to manufacturer’s instructions and residual biotin was removed using a 50kDa MWCO protein concentrator.

Biotinylated pIgG, mAbs and Fabs were titrated using the ELISA method described above. Secondary IgG(H+L) was used for pIgG and mAbs, secondary IgG-kappa was used for Fabs. Maximum binding capacity (1xBmax) was determined, and corresponding concentration was calculated by extrapolation. This way, 0.1xBmax and 10xBmax can also be determined with respective concentrations.

To transfer Abs, desired concentration (0.1xBmax, 1xBmax or 10xBmax) was mixed in PBS to 200µl and were transferred i.p. into recipients. 4 hours post transfer (4hpt), animals were challenged as indicated. 14dpv, animals were sacrificed. Depending on challenging site, spleen or popliteal lymph node (pLN) and blood for serum were harvested. Tissue was processed for flow cytometric staining and serum was used for ELISA.

### Adoptive MBC transfer

C57BL/6 donors were infected with 25 TCID50 PR8, PepBoyIgha recipients with 100 TCID50 J1. At 27dpi, recipients were bled from Vena Saphena. Donors were sacrificed at 28dpi, spleen and medLN were harvested. Tissue was disrupted through a 70µm mesh filter and cells were washed. MBC were isolated using the EasySep™ Mouse B Cell Isolation Kit (Stemcell) according to manufacturer’s instructions, with exception of adding 5µg/sample of anti-IgD biotinylated and anti-GL7 biotinylated mAbs (both Biolegend) to the Abs mix and using 10µl of extra magnetic spheres. After a final wash, isolated MBC are counted using Countess™ 3 Automated Cell Counter (Thermo Fisher) and staining with Acridine Orange (abcam). Some volume of isolated MBC was saved for isolation purity check via flow cytometry.

Priming was necessary to create an environment that supports a robust recall response as transferred MBC did not survive (Supplementary Figure 6A), regardless of CD4 transfer. Additionally, no *wt* HA-specific donor IgG2a (Igh[b]) were detected in naïve mice receiving MBC (Supplementary Figure 6B). Primed J1 mice showed an Ab response that was specific for J1 virus but not for *wt* HA, with only ∼10% of mice having a very low but above background response (Supplementary Figure 6C).

1-5×10^6^ MBC were injected intravenously (i.v.) into recipients in the tail vein. pIgG was transferred at 1xBmax to 200ul with PBS i.p. simultaneously. Animals were left to rest o/n and challenged with UV inactivated virus at 2000 HAU in 100ul i.p. At 7 days post vaccination /8 days post transfer they were sacrificed and spleen, medLN and blood for serum were harvested. Tissue was processed for flow cytometric staining and serum was used for ELISA. For ELISA, secondary biotinylated mAbs IgG2a[a], clone 8.3, and IgG2a[b], clone 5.7, and avidin-HRP (Thermo Fisher) were used. We only considered successful transfers those where at least 50 *wt* HA specific donor B cells (IgM MBC+ switched MBC+ASC+ GC B cells) could be detected after challenge.

### Statistics

Statistic calculations (t-tests, one-way and two-way ANOVA) were performed using the GraphPad Prism Software V.10. Asterisks correspond to: ∗p < 0.05; ∗∗p < 0.01; ∗∗∗∗p < 0.0001 in all figures. All data are represented as mean ± SD.

