## Supplementary Figure for "Antibodies, Memory B Cells, and Antigen Valency Reshape B Cell Responses to Drifted Influenza Virus Vaccination"

### SUPPLEMENTARY FIGURES:

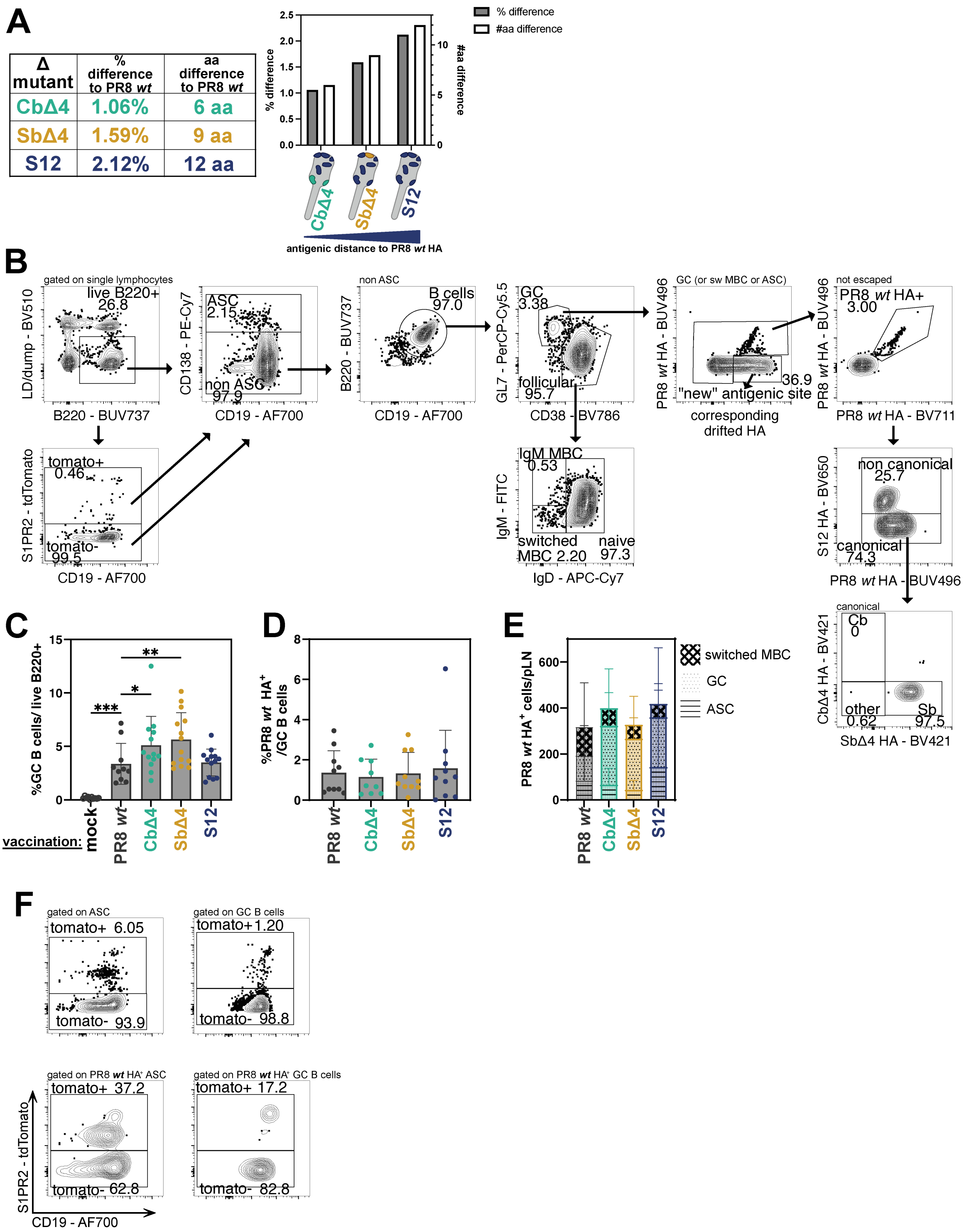


**Supplementary Figure 1: A drifted rHA challenge reshapes secondary GC responses in challenge draining LN. A**: Antigenic distance of CbΔ4, SbΔ4 and S12 to PR8 in % and aa. **B:** Gating strategy used to define PR8 *wt* HA binding of GC B cells, ASC and switched MBC. **C:** Percentage of GC B cells per live B220$^+$, **D:** of PR8 *wt* HA$^+$ GC B cells per GC B cells **E:** and total count of PR8 *wt* HA$^+$ cells among GC B cells, ASC and switched MBC in pLN at 7 days post vaccination (dpv). **F:** Representative flow plots identifying fate mapped (S1pr2-TdTomato$^+$) cells among GC B cells, ASC (top) and PR8 wt HA+ GC B cells and ASC (bottom)

at 7 dpv in pLN. p-values were calculated by one-way ANOVA, with all groups compared to PR8 *wt* HA vaccination (homologous) (C, D). (∗p < 0.05; ∗∗p < 0.01; ∗∗∗∗p < 0.0001). Figures represent data from two experiments with 3-7 mice per challenge group per experiment. At least 2 experiments per group conducted. All data shown as mean ± SD. Animals without any detectable anti-*wt* PR8 HA IgG serum titer at -1 dpv were excluded from data analysis.

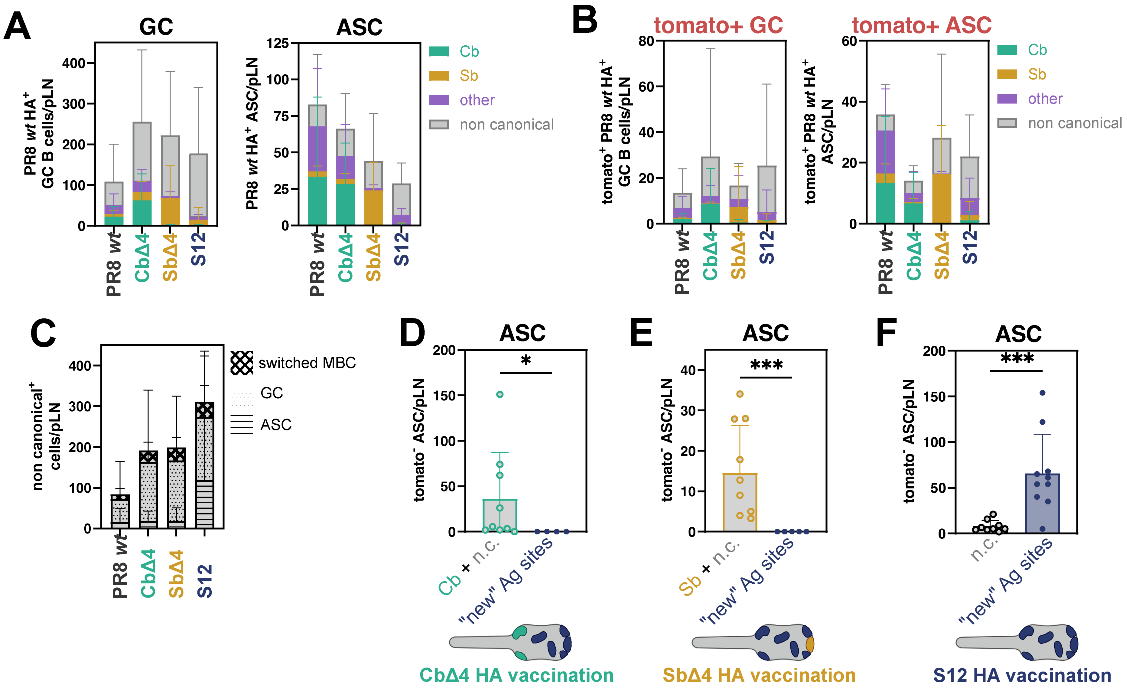


**Supplementary Figure 2. A drifted rHA challenge reshapes immunodominance patterns in draining LN. A:** Immunodominance patterns in Tomato$^-$ and **B:** Tomato$^+$ GC and ASC PR8 *wt* HA$^+$ cells. Specificity to antigenic site indicated by bar color. **C:** n.c. specific cells among GC B cells, ASC and switched MBC at 7 days post vaccination (dpv) in pLN. **D:** Tomato$^-$ Cb$^+$ + n.c.$^+$ and "new" antigenic site$^+$ (PR8 *wt* HA$^-$ CbΔ4 HA$^+$), **E:** Tomato$^-$ Sb$^+$ + n.c.$^+$ and "new" antigenic site$^+$ (PR8 *wt* HA$^-$ SbΔ4 HA$^+$), **F:** Tomato$^-$ n.c.$^+$ and "new" antigenic site$^+$ (PR8 *wt* HA$^-$ S12 HA$^+$) of ASC total count per pLN. p-values were determined by unpaired t-test (**D, E, F**). (∗p < 0.05; ∗∗∗p < 0.001). Figures represent data from two experiments with 3-7 mice per challenge group per experiment. At least two experiments per group conducted. All data shown as mean ± SD. Animals without detectable anti-*wt* PR8 HA IgG serum titers at -1 dpv were excluded from data analysis.

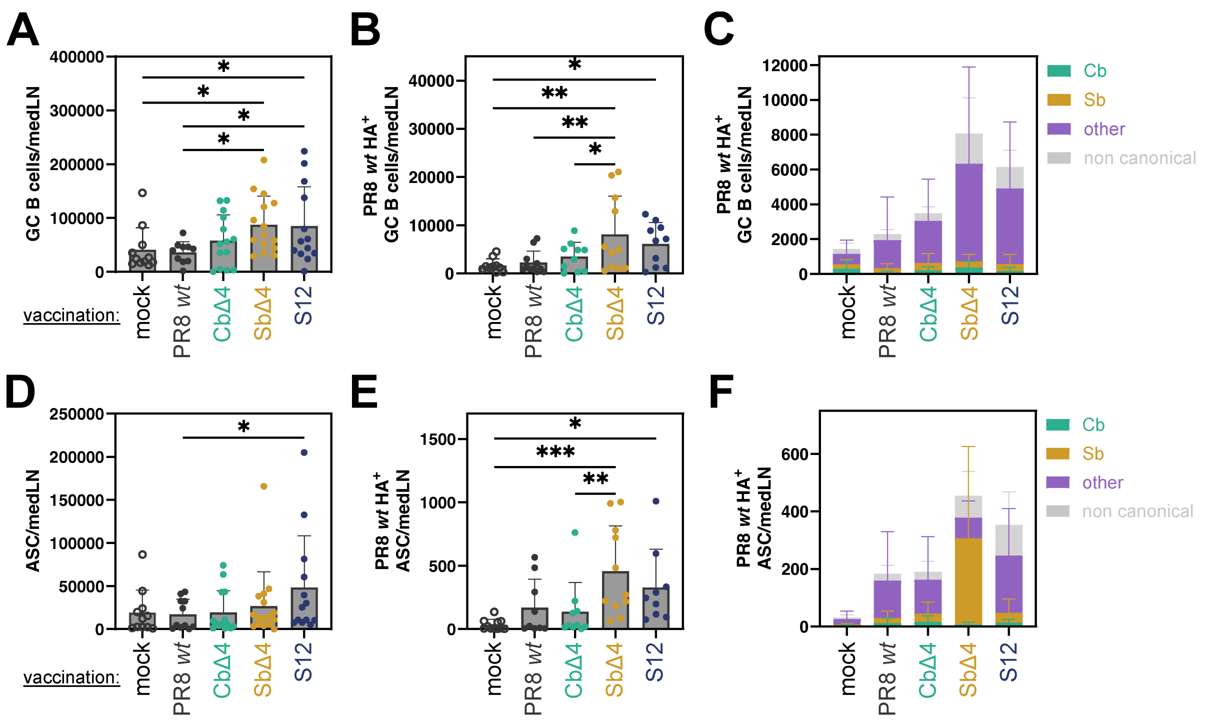


**Supplementary Figure 3: A heterologous rHA challenge reshapes ongoing GC response and immunodominance patterns in prime draining LN.** Total **A:** GC B cells, **B:** PR8 *wt* HA$^+$ GC B cells, and **C:** immunodominance pattern among PR8 *wt* HA$^+$ GC B cells per medLN at 7 days post vaccination (dpv). Total **D:** ASC, **E:** PR8 *wt* HA$^+$ ASC, and **F:** immunodominance pattern among PR8 *wt* HA$^+$ ASC per medLN at 7 dpv. p-values were calculated using by two-way ANOVA (**A, B, D, E**). (*p < 0.05; **p < 0.01; ****p < 0.0001). Lowest p value indicated if no significance detected. Figures represent data from two experiments with 3-7 mice per challenge group per experiment. At least two experiments conducted per group. All data as shown as mean ± SD. Animals without detectable anti-*wt* PR8 HA IgG serum titers at -1dpv were excluded from data analysis.

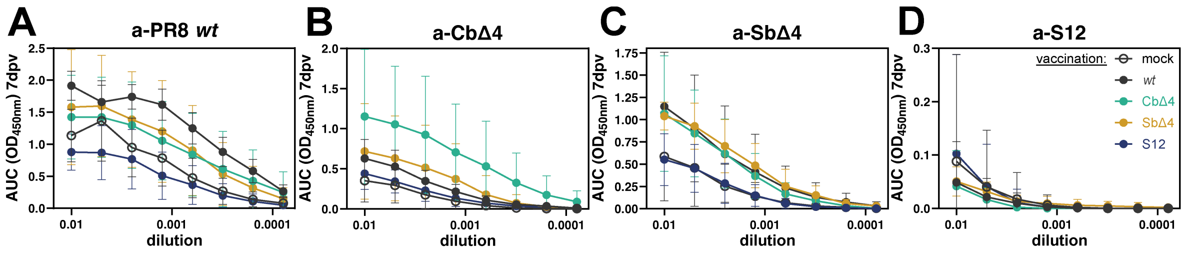


**Supplementary Figure 4: Immunodominance of serum Abs is influenced by drift of vaccine antigen.** OD at 450nm in serum 7 days post vaccination (dpv) after diluation is shown. Plates were coated, from left to right, with rHA: **A:** PR8 *wt*, **B:** CbΔ4, **C:** SbΔ4, **D:** S12. Figures represent data from two experiments with 3-7 mice per challenge group per experiment. All data shown as mean ± SD. Animals without detectable anti-*wt* HA PR8 IgG serum titers at -1dpv were excluded from data analysis.

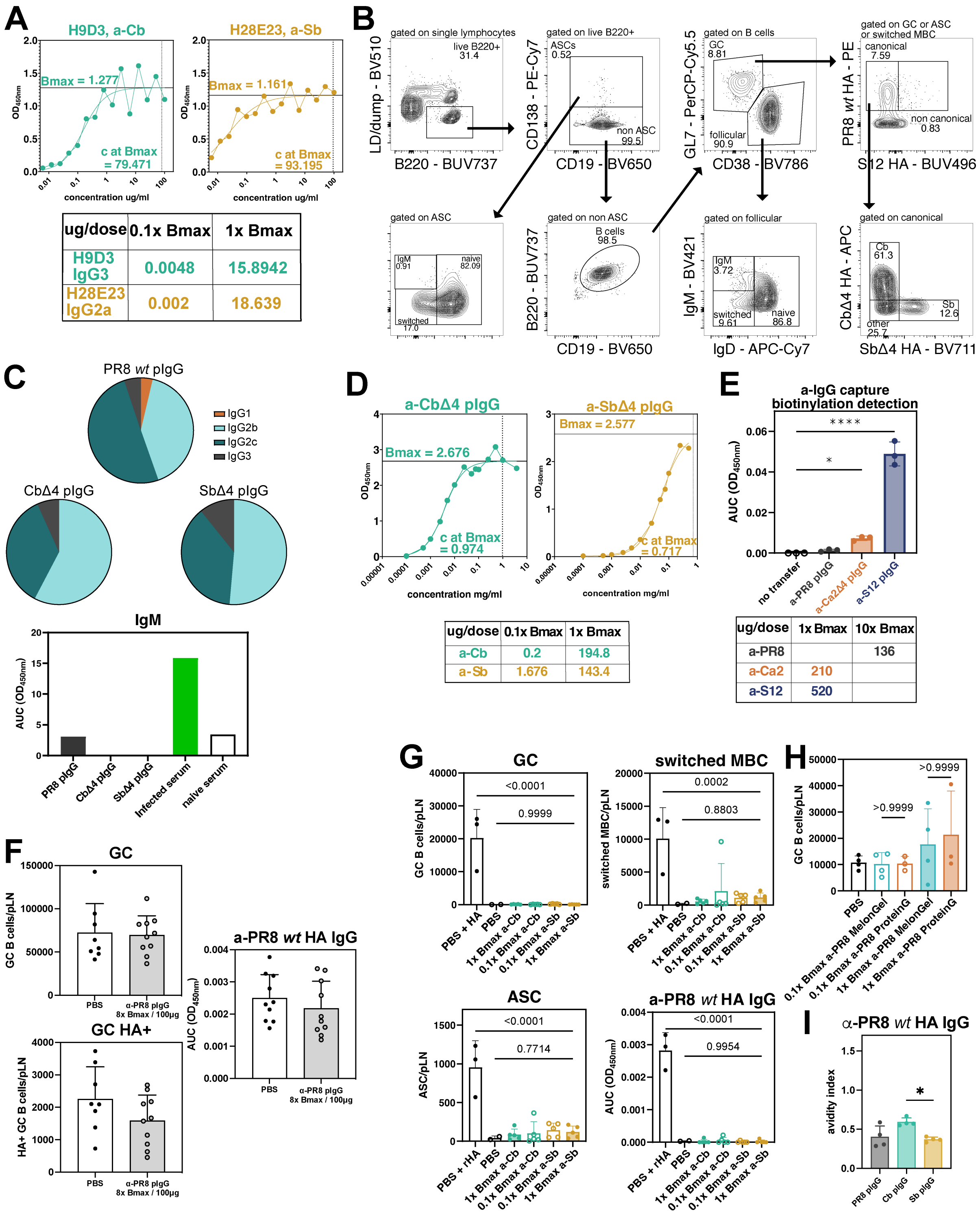


**Supplementary Figure 5: Production and characterization of mAbs and pIgG used for adoptive Ab transfer. A:** Titration and doses of H9D3 and H28E23 at 1x and 0.1x Bmax were calculated using titration. **B:** Gating is shown after flow cytometry at 14 days post vaccination (dpv) of pLN. **C:** Pie charts demonstrating the fractions of IgG subclasses in pIgG mixes (top), and levels of IgM based on AUC OD450nm curves. As control, the total Ig amounts of infected, not processed serum is used (bottom). **D:** Titration and doses of anti(a)-Cb and a-Sb pIgG mixes at 1xBmax and 0.1xBmax were calculated. **E:** Detection of biotinylated pIgG in recipient serum at 4 hours post transfer with goat anti-mouse IgG coated plates and doses of pIgG mixes used. **F:** Total GC B cells, PR8 *wt* $HA^+$ GC B cells and AUC OD450nm of sera at 14dpv after transfer of 8x Bmax a-PR8 pIgG. PR8 *wt* HA was used for coating the plates. **G:** Total cell numbers of ASC, GC B cells and switched MBC per pLN after pIgG transfer without vaccination at d14 (top row). AUC OD450nm of sera after pIgG transfer without vaccination at d14 (bottom row). **H:** Comparison of total GC B cells per pLN at 14dpv after transfer of *wt* HA pIgG purified on a MelonGel column or ProteinG column. **I**: Avidity Index of all pIgG preparations at 1xBmax as determined by a-*wt* HA PR8 ELISA for IgG from OD450nm values. Bmax was determined by nonlinear fit and X (concentration) values were interpolated. p-values were calculated by one-way ANOVA (**I**) with all groups compared to PBS/mock transfer (**E, G**), and MelonGel vs. ProteinG (**H**) shows calculations based on t-tests. (∗p < 0.05; ∗∗p < 0.01; ∗∗∗∗p < 0.0001). Figures represent data from two experiments with 2-3 mice per transfer group per experiment. At least n=2 experiments conducted per group. All data shown as mean ± SD.

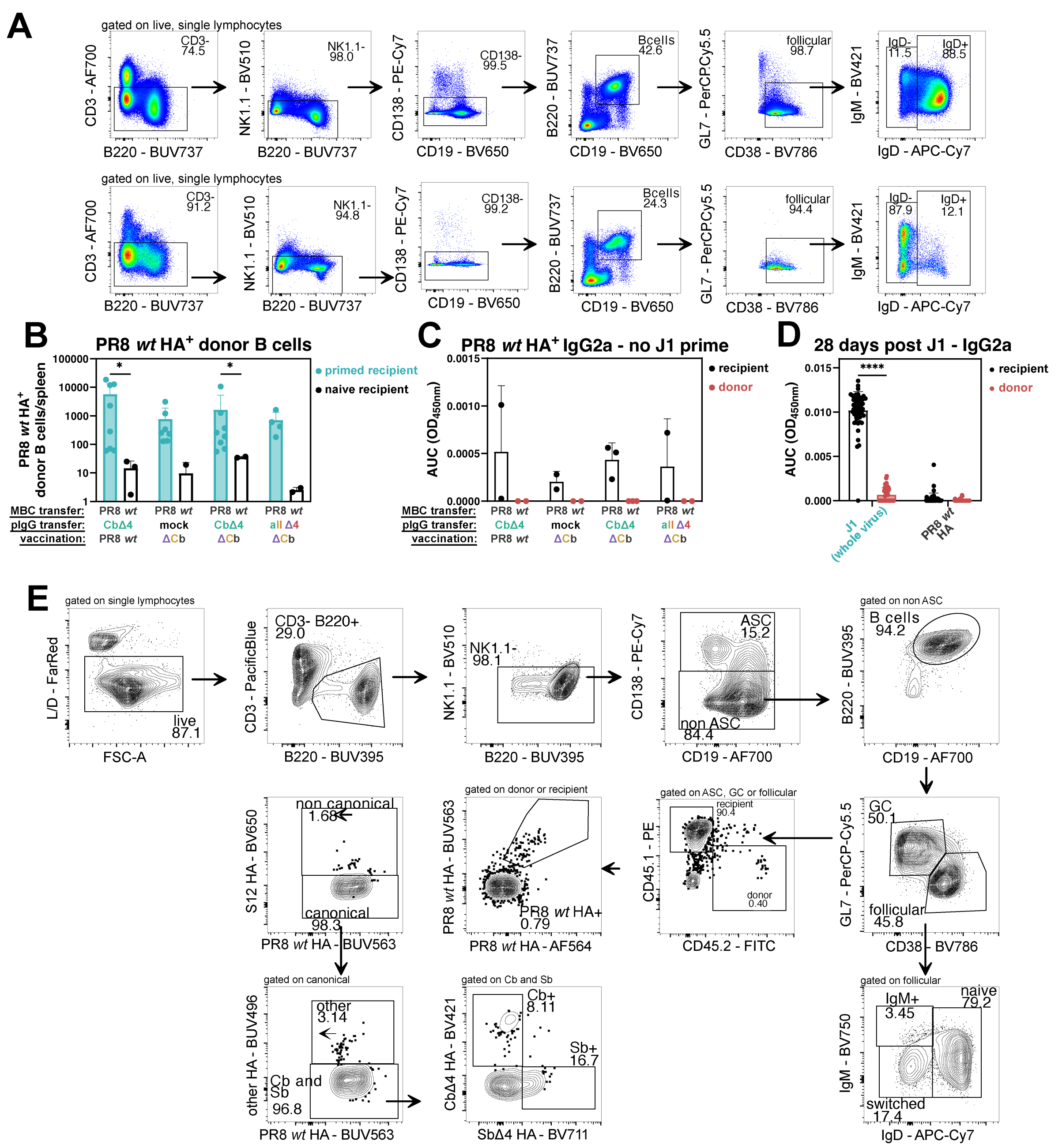


**Supplementary Figure 6: Characterization of donor and recipient responses after MBC transfer. A:** Representative FACS pots of pre- and post-isolation samples. **B:** Total numbers of donor PR8 *wt* HA$^+$ B cells (ASC+GC B cells+IgM MBC+switched MBC) per spleen in naïve and primed recipient are shown. **C:** AUC OD450nm at 7 days post vaccination (dpv) of recipient (Igh[b]) and donor (Igh[a]) PR8 *wt* HA specific IgG2a in naïve donor. **D:** AUC OD450nm at -1dpv from recipient serum against J1 and PR8 *wt* HA for recipient (Igh[b]) and donor (Igh[a]) IgG2a. **E:** Gating strategy for medLN and spleen samples at 7dpv. p-values were determined by unpaired t test (**A, C**). (*p < 0.05; **p < 0.01; ****p < 0.0001). Figures represent data from n=2 experiments/group with 3-7 mice per challenge group per experiment. All data as mean ± SD. Animals with ≤50 donor PR8 *wt* $HA^+$ B cells (ASC+IgM MBC+switched MBC+ GC B cells) per spleen were excluded from analysis.

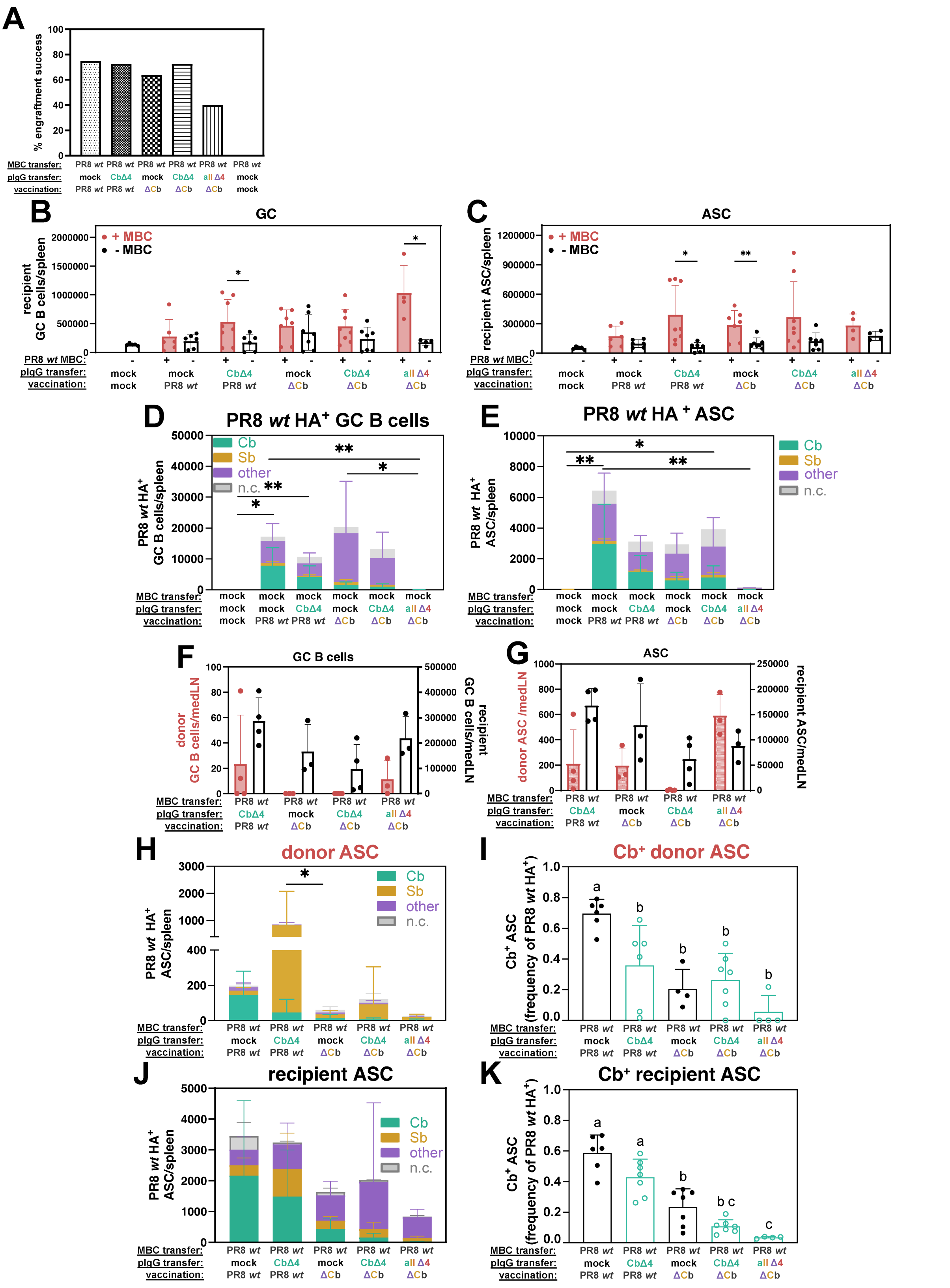


**Supplementary Figure 7: Differential effect of antibodies on naïve B cells and MBC fate and immunodominance and MBC-mediated suppression of antigen-specific naïve B cells.** **A**: Engraftment success rate are shown among all groups tested according to criteria defined. **B:** Total count of recipient GC B cells in spleen and **C:** recipient ASC in spleen with and without MBC transferred are shown. **D:** Immunodominance patterns of recipient PR8 *wt* HA$^{+}$

GC B cells and **E:** ASC in spleen at 7 days post vaccination (dpv) without MBC transfer. **F**: Total count of donor and recipient GC B cells in medLN, **G**: ASC in medLN. **H:** Immunodominance patterns of donor ASC and **I:** frequency of Cb-specific ASC, among PR8 *wt* $HA^+$ cells in spleen at 7 days post vaccination**. J:** immunodominance patterns of recipient ASC and **K:** frequency of Cb-specific ASC, among PR8 *wt* $HA^+$ cells in spleen at 7dpv. p-values were determined by one-way ANOVA (**D, E, H, I, J, K**) or unpaired t test (B, C). ($*p < 0.05$; $**p < 0.01$). For **I** and **K** statistical differences are indicated by compact letter display (cld), where groups with different letters are statistically different. Figures represent data from n=2 experiments/group with 3-7 mice per challenge group per experiment. All data as mean ± SD. Animals with ≤50 donor PR8 *wt* HA+ B cells (ASC+IgM MBC+switched MBC+ GC B cells) per spleen were excluded from analysis.
